# Probe-dependent Proximity Profiling (ProPPr) Uncovers Similarities and Differences in Phospho-Tau-Associated Proteomes Between Tauopathies

**DOI:** 10.1101/2024.03.25.585597

**Authors:** Dmytro Morderer, Melissa C. Wren, Feilin Liu, Naomi Kouri, Anastasiia Maistrenko, Bilal Khalil, Nora Pobitzer, Michelle Salemi, Brett S. Phinney, Dennis W. Dickson, Melissa E. Murray, Wilfried Rossoll

**Affiliations:** Department of Neuroscience, Mayo Clinic, Jacksonville, FL, USA; Proteomics Core, University of California Davis, Davis, CA, USA

## Abstract

Tauopathies represent a diverse group of neurodegenerative disorders characterized by the abnormal aggregation of the microtubule-associated protein tau. Despite extensive research, the precise mechanisms underlying the complexity of different types of tau pathology remain incompletely understood. Here we describe an approach for proteomic profiling of aggregate-associated proteomes on slides with formalin-fixed, paraffin-embedded (FFPE) tissue that utilizes proximity labelling upon high preservation of aggregate morphology, which permits the profiling of pathological aggregates regardless of their size. To comprehensively investigate the common and unique protein interactors associated with the variety of tau lesions present across different human tauopathies, Alzheimer’s disease (AD), corticobasal degeneration (CBD), Pick’s disease (PiD), and progressive supranuclear palsy (PSP), were selected to represent the major tauopathy diseases. Implementation of our widely applicable Probe-dependent Proximity Profiling (ProPPr) strategy, using the AT8 antibody, permitted identification and quantification of proteins associated with phospho-tau lesions in well-characterized human post-mortem tissue. The analysis revealed both common and disease-specific proteins associated with phospho-tau aggregates, highlighting potential targets for therapeutic intervention and biomarker development. Candidate validation through high-resolution co-immunofluorescence of distinct aggregates across disease and control cases, confirmed the association of retromer complex protein VPS35 with phospho-tau lesions across the studied tauopathies. Furthermore, we discovered disease-specific associations of proteins including ferritin light chain (FTL) and the neuropeptide precursor VGF within distinct pathological lesions. Notably, examination of FTL-positive microglia in CBD astrocytic plaques indicate a potential role for microglial involvement in the pathogenesis of these tau lesions. Our findings provide valuable insights into the proteomic landscape of tauopathies, shedding light on the molecular mechanisms underlying tau pathology. This first comprehensive characterization of tau-associated proteomes across different tauopathies enhances our understanding of disease heterogeneity and provides a resource for future functional investigation, as well as development of targeted therapies and diagnostic biomarkers.

## Introduction

Abnormal accumulation of the microtubule-associated protein tau as insoluble fibrils, is the major neuropathologic hallmark precipitating neuronal and glial cell death within a group of clinically [111, 121], biochemically [154] and pathologically [21] heterogenous neurodegenerative disorders, termed tauopathies [86]. Tau is encoded by the *MAPT* gene located on chromosome 17 in the human genome and plays an important role in the organization of microtubules operating within the dynamic cytoskeleton [39, 118]. Under pathological conditions, tau dissociates from microtubules, undergoes misfolding, is subject to extensive post-translational modifications, and forms filamentous aggregates [53, 83]. Immunohistochemical staining of postmortem tau aggregates with phospho-tau specific antibodies, such as AT8, remains the gold-standard for neuropathologic diagnosis [32, 35, 42] (Fig.1). Clinicopathologic and genetic studies have provided significant evidence that age-at-onset, disease duration, severity of antemortem cognitive decline and phenotypic symptoms are driven by the burden of tau lesions [11, 15, 52, 70, 79, 85, 96, 98, 112, 113, 126, 137, 142, 151]. In addition, over 60 genetic mutations in tau have been linked to neurodegenerative diseases [62, 63, 144]. These findings highlight the critical importance of developing therapeutic strategies targeting modifiers as well as differentially diagnostic or early-detection biomarkers of tau pathology [8, 23, 50, 60, 76, 78, 159].

**Fig. 1.**
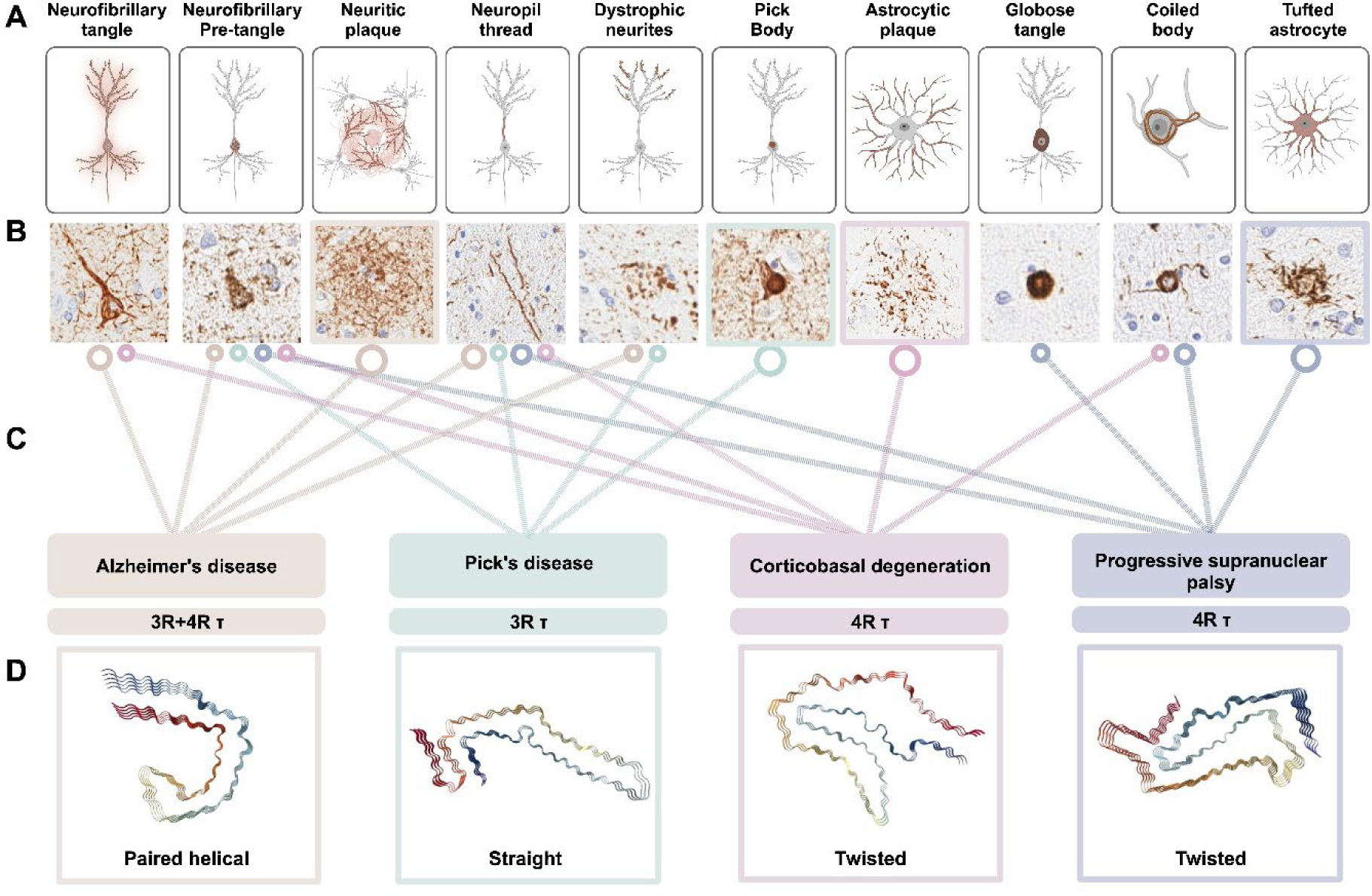
Neuropathological and structure-based characteristics of major tauopathies used for in situ Probe-Dependent Proximity Proteomic profiling of the phospho-tau associated proteome. **(A)** Schematic depicting the cellular localization of distinct and shared phospho-tau aggregates across Alzheimer’s disease (AD), Pick’s disease (PID), corticobasal degeneration (CBD) and progressive supranuclear palsy (PSP). **(B)** Representative human neuropathology images of AT8 stained human brain tissue. **(C)** Lines indicating the presence of each type of neuropathological lesion in different tauopathies, size of circular end symbol denoting the presence of a defining pathological hallmark specific to either mixed 3- and 4-repeat, 3-repeat or 4-repeat tauopathies. **(D)** CryoEM filament structure images generated from RCSB Protein Database (PDB) using the NGL viewer [134] of **PDB-ID 503L** (AD, *Fitzpatrick et al., 2017* [49], PDB https://doi.org/10.2210/pdb5O3L/pdb), **PDB-ID 6GX5** (PiD, *Falcon et al., 2018* [46] PDB https://doi.org/10.2210/pdb6GX5/pdb), **PDB-ID 6JTO** (CBD, *Zhang et al., 2020*, [177] PDB https://doi.org/10.2210/pdb6TJO/pdb) and **PDB-ID 7P65** (PSP, *Shi et al., 2021* [147] PDB https://doi.org/10.2210/pdb7P65/pdb).

To date, a spectrum of over twenty-six diverse pathological phenotypes have been described [86, 143]. Tauopathies are classified as primary or secondary taxonomies, depending on whether tau aggregates are the sole pathologic cause or exist within a multi-proteinopathy, respectively [3]. The most common secondary tauopathy is Alzheimer’s disease (AD), where Thal Phase denotes the stereotypic spread of extracellular amyloid-beta (Aβ) plaque pathology [158], while Braak stage defines tau pathology [14]. AD tau hallmarks include neurofibrillary tangles (NFTs), neuritic plaques (NPs), neuropil threads (NTs), and tangle associated neuritic clusters (TANCs) [108]. Frontotemporal lobar degeneration with tau pathology (FTLD-tau) includes the primary tauopathies such as Pick’s disease (PiD), corticobasal degeneration (CBD), and progressive supranuclear palsy (PSP) [21]. PiD is a rare disease that is defined by its unique accumulation of spherical intraneuronal argyrophilic inclusions of Pick bodies (PiBs), accompanied by ballooned neurons (BNs/Pick cells), dystrophic neurites (DNs) and small glial inclusions [34, 65]. PSP is histopathologically diagnosed based on the exclusive presence of tufted astrocyte (TA) lesions in pre-defined brain areas [133], accompanied by globose ballooned neurons (GLO/BN) and premature neurofibrillary tangles (PTs), oligodendroglial coiled bodies (CBs) and neuropil threads (NTs) [36, 120]. CBD is the most common primary tauopathy [121], and shares a clinico-pathological spectrum of symptom presentation and mutual pathologic inclusions with PSP [152], yet is characterized by the distinctive presence of astrocytic plaques (AP), alongside CBs, NFTs and NTs [82]. Primary age-related tauopathy (PART) is a discrete tauopathy characterized by NFT tau pathology restricted mostly to the medial temporal lobe, associated commonly with aging, and not associated with concomitant temporal Aβ pathology [26].

The neuropathological classification of these tauopathies is supported by the biochemical and structural nature of the tau proteoforms present in disease aggregates, namely the splicing isoforms containing three or four microtubule-binding repeats (3R or 4R tau) [53] and more recently, structure-based classification of tau filament folds [147] (Fig. 1). Primary tauopathies exclusively formed by 4R tau isoforms into inclusions include CBD and PSP, 3R tau isoforms are primarily affiliated with PiD pathology, whereas AD and PART are mixed 3R/4R tauopathies [21]. Cryogenic electron microscopy (cryo-EM) studies provide <6 Å super-resolution maps of filament periodicity and atomic modeling of fold structure, main chain conformation and inter-protofilament interfaces [147]. The discovery of tauopathy-specific ultra-structural taxonomy of tau protofilaments and conformational folds confirms disease-specific biochemical diversity among this group of disorders [2, 46–49, 147, 177]. The potential of tau aggregates to differentially seed formation of structurally analogous inclusions in recipient cells suggests that the heterogenous clinical and pathological manifestation of tauopathies can be defined by distinct tau strains [72, 138, 155, 164]. [2] It remains critically important to identify common and disease-specific tau pathology-associated proteins that act as modifiers of tau aggregation and spread, as mediators of cellular pathologies, and as tauopathy-specific biomarkers.

Profiling of tau-associated proteomes through proximity proteomics and immunoprecipitation-based interactome studies have aimed to identify modifiers and pathways altered during disease pathogenesis, in a variety of cellular and animal model systems [73]. In addition, profiling the proteome of large NFT somal inclusions in AD has been performed via laser-capture microdissection previously [41]. However, the proteomes associated with the entire burden of distinct tau lesions have not been systematically assessed across different diseases within the same study. Specifically, proteomic analysis of neuropil thread pathology and the finer axonal and dendritic extensions of neuronal and glial tau aggregates have not yet been performed, which require inclusion within proteomic profiling studies. Major limitations for previous reports include lack of disease modelling of human-distinct tau strains, making post-mortem human tissue the only viable source of studying heterogeneous disease-relevant tau aggregates. Therefore, the analysis of morphologically well-preserved pathology, exhibiting disease relevant protofilament structure of tau isoforms, with cell type specific and region-dependent vulnerability, has the potential to reveal novel protein interactors and disease modifiers. Several groups have initiated utilization of biochemical methodology to probe target proteins *in situ*, based on rapid biotin-tyramide labelling of proximal proteins, mediated by horseradish peroxidase (HRP) conjugated to a primary antibody against the target protein. This approach is based on enzyme-mediated activation of radical sources (EMARS) [56, 81] and has been termed as Selective Proteomic Proximity Labeling Assay Using Tyramide (SPPLAT) for the proteomic analysis of membrane-bound protein clusters in cell culture [88, 130] and subsequently as Biotinylation by Antibody Recognition (BAR) for labelling proteins associated with lamin A/C in tissue [7, 130]. Since then this approach has also been used to profile the associated proteome of alpha-synuclein aggregates in Parkinson’s disease and Lewy Body Dementia [75], and, more recently, in a proof-of-concept workflow to identify the phospho-tau associated proteome in PSP, yet lacking immunohistochemical (IHC) candidate validation [129]. Here, we have established a related strategy with important technical modifications under the suggested designation of <u>Pro</u>be-dependent <u>P</u>roximity <u>Pr</u>ofiling (ProPPr), to reflect its extended flexibility with regard to target-specific probes (e.g antibodies, oligos, toxins, aptamers etc.) and the nature of target-proximal molecules (proteins or nucleic acids) that can be labelled. In the current study, we have used ProPPr with the AT8 antibody for the unbiased profiling of the phospho-tau associated proteomes across the 4 major tauopathies (AD, CBD, PiD, and PSP), to quantify both shared and disease-specific proteins associated with these heterogenous protein aggregates. Subsequent IHC validation of selected proximal proteins in post-mortem brain tissue within specific subsets of tau lesions, highlights their implications in tauopathies for further research into novel therapeutic strategies and biomarkers.

## Methods

### Human post-mortem case selection and neuropathologic evaluation

A cohort of human post-mortem cases were selected from brains donated for research to the Mayo Clinic Florida brain bank for neurodegenerative disorders. The brain bank accommodates cases from the Mayo Clinic Alzheimer’s Disease Research Center, the Mayo Clinic Study of Aging, the Einstein Aging Study, CurePSP, and the State of Florida Alzheimer’s Disease Initiative. Informed consent was obtained for brains donated for research studies and samples were de-identified, as approved by the Mayo Clinic Institutional Review Board for the purpose of autopsy research. De-identified studies of autopsy samples are considered exempt from human subject research regulations. Core demographic and clinical data to generate the cohort were collected from available clinical history notes and the Mayo Clinic Florida Brain Bank database. Age at onset (AAO) detailed the time that either the patient or caregiver first reported symptoms of impairment. Disease duration was calculated as the interval period between AAO and age at death. Sex and ethnoracial status were self-reported by patients. Brains were processed in 10% neutral buffered formalin followed by systematic sampling of tissue after gross neuroanatomical examination of the fixed specimen, where brain weight was calculated from the fixed hemibrain and doubled to obtain total brain weight (g). Cortical sections were sampled perpendicular to the gyrus to maintain integrity of the full cortical ribbon. Blocked tissue samples were retained in cassettes in 10% formalin prior to preservation and sectioning in paraffin.

Histological assessment for post-mortem diagnosis was conducted on 5 µm formalin-fixed paraffin embedded (FFPE) tissue sections mounted on positively charged glass slides by a single expert neuropathologist (D.W.D), following standardized neuropathologic methods for complete characterization of cases, as previously described [139]. Briefly, cases were examined in line with established guidelines for the criteria to satisfy the diagnosis of intermediate to high likelihood of AD (CERAD, [106]; NIA-AA, [64, 102, 109, 110]), FTLD-tau [4, 16, 57, 133] or PART [26]. Cerebrovascular disease [25, 31, 111, 157], alpha-synuclein [101, 162], and TDP-43 pathology [71, 95, 100, 116, 117] was also examined for presence of concomitant neuropathologic change in accordance with recognized guidelines.

Tissue blocks corresponding to frontal cortex (middle frontal gyrus, Brodmann area 9) were used for proximity labelling studies. The choice of this area met the requirements for study design to use a cortical brain region that was shared amongst tauopathy groups and demonstrated disease-specific tau pathology, whilst retaining equal background of cell type and molecular tissue composition. The disease cohort in this study was curated to include the four major tauopathy disorders (AD, CBD, PiD, PSP) with pre-defined inclusion/exclusion criteria. AD cases were selected with significant pathology reaching Braak tangle stage >IV [13, 14], Thal amyloid phase >4 [158], and high probability AD according to current recommended National Institute on Aging (NIA)-Alzheimer’s Association and NIA-Reagan criteria [1, 64, 102, 109, 110]. PSP and CBD cases were identified according to Rainwater and FTLD-tau criteria [4, 16, 57, 133]. The presence of cerebrovascular disease was evaluated using a 10-point modified Kalaria score [31], and cases were excluded based on a Kalaria score ≥4 for the cortex and basal ganglia regions assessed. Significant coexisting neuropathologic changes provided additional exclusion criteria, including hippocampal sclerosis [115], presence of TDP-43 [71, 95, 100, 116, 117] or α-synuclein pathology [101, 162] in the amygdala, brainstem, or cortex. Exclusive criteria were also applied to aged controls displaying the common and mild aging associated process of PART, these cases were included for comparison that have little to absent AD pathology in BA9 (Braak stage ≤ III, Thal phase < 1), had no other co-pathologies [31, 95, 100, 115] and lacked cognitive decline in life [26]. The curation of frontal cortex permits the use of PART cases without frontal pathology in the context of this commonly observed aging process. Demographic and clinicopathologic data of all patient cases used in the study are provided in the Supplementary Table 1.

### Quantitative digital pathology

The first 5 µm FFPE tissue section from the series of slides for each case later used for AT8 ProPPr, were immunostained on a Thermo Fisher Lab Vision 480S autostainer by the Mayo Clinic Florida Brain Bank Core. Immunohistochemical staining was performed using phospho-tau antibody (AT8, 1:1000, phospho-S202/T205, Thermo Fisher Scientific, MN1020) and diaminobenzidine detection with hematoxylin counterstain. Digital pathology methods using Leica’s Aperio technology have been previously described [107, 111]. In brief, AT8 and hematoxylin-stained slides were digitized using the Aperio AT2 scanner, image files were traced and annotated using Aperio ImageScope software (Leica Biosystems, version 12.4.3.5008). Batch analysis was conducted in eSlideManager (Leica), where color deconvolution and custom macros measured phospho-tau burden on AT8 stained sections to produce quantitative pathology data for each case (% tau burden) (Supplementary Table 1). Representative high-resolution images were acquired at 2x and 20x using ImageScope and exported for demonstrative purposes (Fig.1 - Fig.2).

**Fig. 2.**
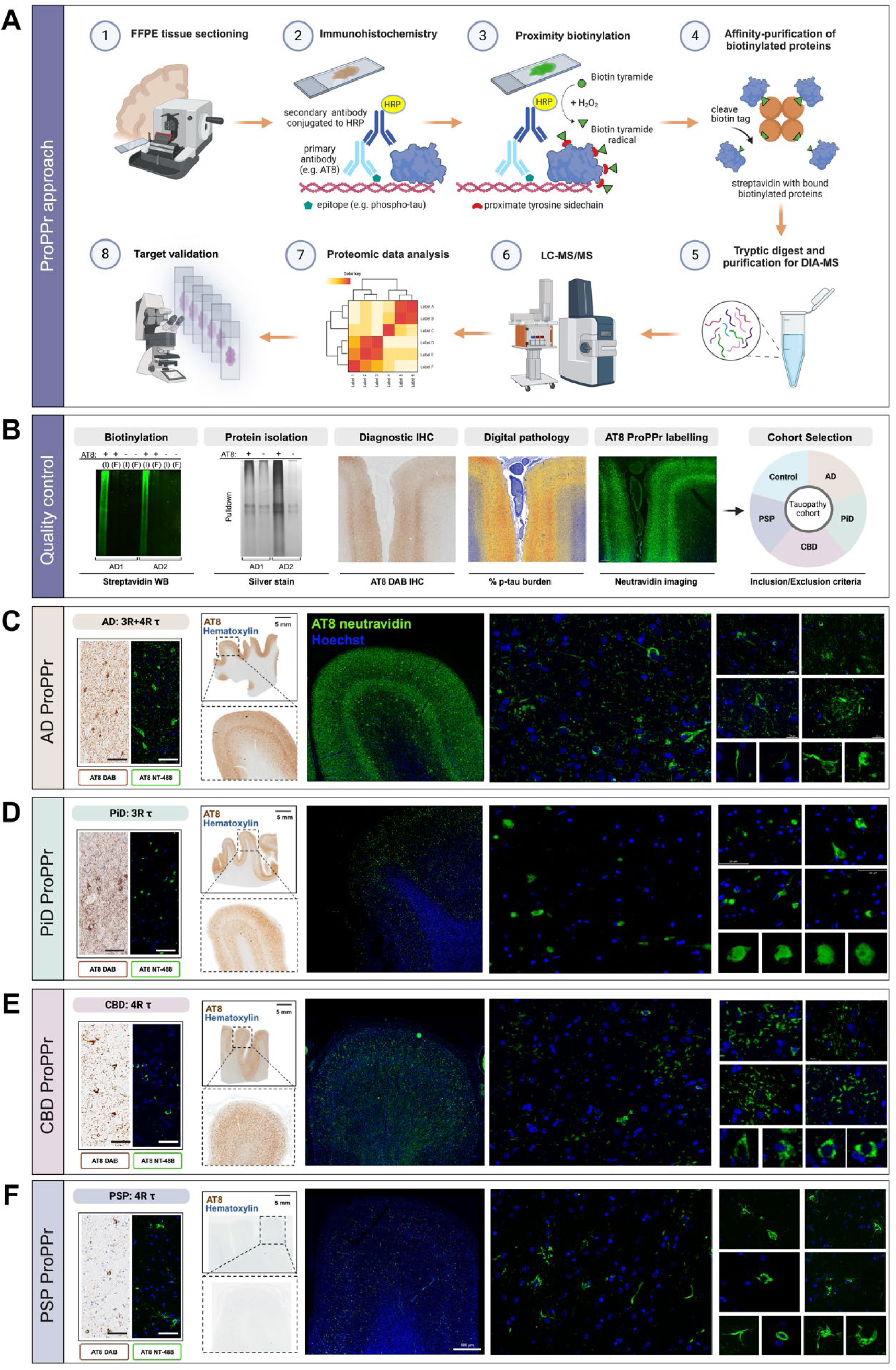
Validation of Probe-dependent Proximity Profiling (ProPPr) method. **(A)** Schematic representation of the ProPPr method. **(B)** Western blotting of the tissue lysates reveals extensive biotinylation in the AD samples treated by ProPPr method, but not in the negative controls where staining with AT8 antibodies was omitted. I – input lysate; F – flow-through after pulldown with streptavidin beads. Silver staining of the proteins eluted from streptavidin beads after pulldown reveal increased level of protein in the ProPPr samples comparing to negative controls where staining with AT8 antibodies was omitted. AT8 ProPPr quality control images compared distribution, tau burden by digital pathology and neutravidin labelling of AD frontal cortex. **(D-F)** Representative images of routine immunohistochemistry and fluorescent proximity-biotinylation labelling of phospho-tau aggregates by AT8 ProPPr method of major tauopathies, AD **(C)**, PiD **(D)**, CBD **(E)**, and PSP **(F)** show matching disease-specific patterns of phospho-tau pathology, when FFPE tissue from the cortex was processed for routine DAB staining with phospho-tau (AT8) antibody and hematoxylin counterstain and adjacent sections fluorescently labelled using AT8 ProPPr and visualized using NeutrAvidin Dylight 488. Staining patterns on adjacent sections are evenly matched in the immunohistochemistry and proximity-biotinylation of phospho-tau in macroscopic cortical tile images and high magnification microscopy demonstrates the specific labelling of tau lesions found in each tauopathy.

### Phospho-tau proximity labelling

Phospho-tau proximity labelling for proteomics with immunofluorescent controls were performed on all cases using 5 µm FFPE frontal cortex tissue sections. Tissue was deparaffinized by immersion in xylene and rehydrated through a series of graded ethanol solutions (100%, 90% and 70% ethanol). Endogenous peroxidase activity was quenched using immersion in absolute methanol containing 0.3% H_2_O_2_ (Sigma-Aldrich) for 20 min. Antigen retrieval using heated citrate buffer, pH 6 (Dako Target Retrieval Solution, S2369), was induced in a steamer for 30 min. Tissue sections in solution were cooled to room temperature (RT) and washed with a continuous flow of dH_2_O for 10 min. Following tissue rinsing with dH_2_O for 5 minutes, sections were concurrently blocked and permeabilized using 10% normal goat serum (NGS, Sigma-Aldrich) in Tris-buffered saline (TBS, DAKO), containing 0.3% Triton-X 100, at RT for 1 h. Tissues were incubated with AT8 phospho-tau antibody (1:500), in antibody diluent (5% NGS with 0.3% Triton X-100 in TBS) and incubated overnight at 4°C in a humidified chamber. Tissue sections assigned to negative control samples were incubated overnight in antibody diluent without adding primary antibodies. Tissue sections were washed three times (5 min each) in TBS containing 0.05% Tween-20 (TBS-T). Antibody diluent containing secondary antibody (1:500, F’(ab)2 goat anti-mouse cross-adsorbed HRP, Invitrogen, A24524) was incubated at RT for 1 h, followed by three 5 min TBS-T washes. Immunostained sections were washed 3×5 minutes with Wash Buffer followed by HRP-mediated protein proximity labelling in Labelling Buffer (1xTBS, 5 uM biotin-ss-tyramide (Iris Biotech, LS-3570), 0.03% H_2_O_2_) for 5 min. The reaction was stopped by 5-minute incubation of slides in Quenching Buffer (1xTBS, 0.5 M sodium ascorbate).

Fluorescent NeutrAvidin labelling on the control section from each case was used as a quality control to ensure adequate biotin-tyramide labelling of each case used for proteomic mass-spectrometry analysis and to serve as an imaging sample to demonstrate the spatial labelling capacity of pathology by AT8-positive phospho-tau proximity labelling. After biotin-SS-tyramide labelling, the experimental control sections were washed three times (5 min each) with TBS-T. NeutrAvidin DyLight-488 (1:500, Invitrogen, 22832) was diluted in antibody diluent and incubated for 1h at RT. Tissue was washed with TBS-T and incubated with Hoechst 33342 (1:1000, Thermo Fisher Scientific, 62249) for 20 min at before rinsing with TBS. To quench autofluorescence, tissues were stained with 1x TrueBlack lipofuscin autofluorescence quencher (Biotium, 23007) for 30 sec and rinsed with dH_2_O and mounted with glass coverslips using ProLong Glass Antifade Mountant (Invitrogen, P36980).

### Affinity purification of biotinylated proteins

After proximity labelling antibodies were first stripped from the tissue sections by incubation with 25 mM glycine, pH 2.0 for 30 min. Tissue was than scraped from the glass slides with razor blade and placed in 0.5 ml Precellys tubes supplemented with CK14 beads for homogenization. For each sample six 5 um sections were combined in one tube. Tissue was supplemented with 200 uL Lysis Buffer I (50 mM ammonium bicarbonate, 0.2% RapiGest SF (Waters)) and homogenized in Precellys 24 instrument (Bertin Technologies) by applying two 30 s cycles of homogenization at 6500 rpm with 30 s interval between the cycles. Homogenates were transferred to 0.2 uL PCR tubes and incubated at 95 °C for 30 min. Lysates were sonicated for 30 s at 40% amplitude using Qsonica Q125 sonicator and further incubated at 80 °C for 2 hours. Lysates were cleared by centrifugation at 21,000 g for 15 min. Supernatants were collected in clean 1.5 ml Protein LoBind Tubes (Eppendorf), and pellets were resuspended in 200 uL Lysis Buffer II (50 mM ammonium bicarbonate, 8 M urea), incubated for 15 min at RT and centrifuges for 15 min at 21000 g. Supernatants was mixed with the cleared lysates from previous centrifugation. 5% aliquots from the resulting samples were stored for further mass spectrometry analysis of total lysates, and the remaining samples were incubated with 20 uL Streptavidin Sepharose High Performance beads (Cytiva, 17511301) overnight at 4 °C with rotation. Next day beads were washed 2 times in Wash Buffer I (50 mM ammonium bicarbonate, 0.2% Zwittergent 3-16) and 2 times in Wash Buffer II (50 mM ammonium bicarbonate, 6 M urea). Biotinylated proteins were then eluted from the beads by reducing disulfide bond in the linker between biotin and tyramide with Elution Buffer (50 mM ammonium bicarbonate, 6 M urea, 10 mM TCEP) using two 15 min incubations at 37 °C.

### Mass spectrometry

Total lysate samples were reduced with 10 mM TCEP in 50 mM ammonium bicarbonate buffer for 30 min at 37 °C and alkylated with 20 mM solution of iodoacetamide in 50 mM ammonium bicarbonate buffer for 30 min at room temperature in the dark. Proteins eluted from the beads were alkylated the same way without the reduction step. Proteins were then incubated with Lys-C/trypsin mixture (Promega) (200 ng for pull-down samples and 1 ug for total lysate samples) for 2 hours at 37 °C. Afterwards the samples were diluted to decrease the concentration of urea up to 1 M and supplemented with additional amounts of sequencing-grade trypsin (Promega) (200 ng for pull-down samples and 1 ug for total lysate samples), followed by overnight incubation of the samples at 37 °C. Peptides were directly loaded on an Evosep C18 tip and separated using the Evosep One and the 100 spd High Organic method (Evosep, Odense, Denmark). Peptides were eluted and ionized using a Bruker Captive Spray emitter. A Bruker timsTOF Pro 2 mass spectrometer running in diaPASEF mode [103] was used for acquisition. The acquisition scheme used for diaPASEF consisted of 6×3 50 m/z windows per PASEF scan.

### Proteomic data analysis

Data-independent acquisition (DIA) mass spectrometry data was analyzed with the Spectronaut 17 software package (Biognosys) using the targeted library-based approach. Firstly, a spectral library was generated by searching the data for both pull-down and total lysate samples using the Spectronaut pulsar search engine with the default setting against the Uniprot UP000005640 Human database (20,607 entries) supplemented with the common contaminants database (38 entries). Pulldown samples were separately matched with the resulting library using the Spectronaut 17 default settings. Briefly, “trypsin/P specific” was set for the enzyme allowing two missed cleavages, fixed modifications were set to cysteine carbamidomethylation, and variable modification were set to peptide N-terminal acetylation and methionine oxidation. For DIA search identification, PSM and Protein Group decoy false discovery was set at 1%. Protein level intensities were summarized using the MaxLFQ algorithm. Phospho-tau associated proteins were identified using the MiST computational tool for scoring of affinity purification mass spectrometry data using pull-down samples and their corresponding negative controls for each pathology separately [66]. For the analysis of differential protein association with AT8-positive phospho-tau in different pathologies, data normalization, imputation of missing values and differential abundance test was performed using DEP R package (version 1.22.0) [179]. Proteins were considered significantly different in each of the pairwise comparisons when adjusted p-value was less than 0.05. GO term overrepresentation analysis was performed using the clusterProfiler R package (version 4.8.2) [174]. GO term networks were generated by the EnrichmentMap plugin (version 3.1.0) and protein-protein interaction network was imported from the StringDB database using the stringApp plugin (version 2.0.2) in Cytoscape software (version 3.9.1) [37, 105, 146]. Venn diagrams were generated using the online tool for calculation and drawing of Venn diagrams developed in the University of Ghent (https://bioinformatics.psb.ugent.be/webtools/Venn/). The statistical significance of the overlap between gene sets was calculated by hypergeometric test using the online tool from the Graeber lab at the University of California Los Angeles (https://systems.crump.ucla.edu/hypergeometric/index.php). Plots were created in Rstudio development environment (version 2023.06.1) for R programming language (version 4.3.1) using ggplot2 R package [168].

### Immunohistochemical co-immunofluorescence validation of protein candidates

Additional 5 µm FFPE frontal cortex tissue sections were used from select cases for validation of candidate proteins from proximity labelling studies by co-immunofluorescence tissue staining. Tissue was treated as described above for proximity labelling using the methods of deparaffinization, antigen retrieval, whilst omitting endogenous peroxidase quenching, with only immersion in absolute methanol for 20 min. Following tissue rinsing with dH_2_O for 5 minutes, sections were blocked and permeabilized in tandem using serum-free protein block (Dako, S0909) containing 0.3% Triton-X 100 at RT for 1h. Tissues were immunostained with the following primary antibodies for selected candidates and co-staining analysis: AT8: MN1020, Invitrogen (1:600), VPS35: GTX108058, GeneTex (1:400), VGF: ab308287, Abcam (1:200), FTL: NBP2-34072, Novus (1:400) GFAP - 2.2B10, Invitrogen (1:1000), IBA1 - ab300156, Abcam (1:500). Antibodies were diluted in antibody diluent (Dako, S0809) containing 0.3% Triton X-100 and incubated overnight at 4 °C in a humidified chamber. Sections were washed with TBS-T, three times for 10 min each. Alexa Fluor conjugated secondary antibodies were diluted in antibody diluent and incubated for 2 h at RT with the omission of light sources. After incubation sections was washed with TBS-T and incubated with Hoechst 33342 (1:500, Thermo Fisher Scientific, 62249), tissue was stained with tissues were stained with 1x True Black lipofuscin autofluorescence quencher (Biotium, 23007) and mounted as described above for NeutrAvidin proximity labelling sections. High-resolution imaging was conducted, and a series of optical sections were acquired spanning the 5 µm depth of the tissue. Z-series were obtained using an Eclipse Ti2 fluorescence microscope (Nikon) equipped with a Spectra X multi-LED light engine (Lumencor), single bandpass filter cubes for DAPI, EGFP/FITC, TRITC and Cy5 (Chroma), and a Zyla 4.2 PLUS sCMOS camera (Andor), using NIS Elements HC V5.30 software (Nikon). Out of focus light from each z-series was eliminated using the Clarify.ai module of the NIS-Elements Advanced Research (V5.30) deconvolution package. Clarified z-series are shown as optical projections and 3D volume rendered views to demonstrate colocalization of tau pathology and proteomic candidate markers. Fluorescent NeutrAvidin-488 images and triple labelling tile images were captured using z-stacks and full focus projection through a 40x objective on an inverted fluorescent microscope (Keyence BZ-X800) and high-resolution single-plane tile images were stitched using the Keyence analyzer package (BZ-X series). Select NeutrAvidin-488 and candidate validation co-localization was captured using z-stacks at 60x or 100x on a Keyence BZ-X800 using high resolution capture mode, full focused images were created the analyzer package.

### Tau ProPPr candidate validation criteria

Co-immunofluorescence was conducted after optimization of primary antibody dilution and antigen retrieval, the fluorescent secondary antibodies for two markers (i.e. AT8 and candidate protein) used were anti-rabbit F(ab’)2-AlexaFluor-488 and anti-mouse F(ab’)2-AlexaFluor-647 to reduce the possibility of spectral crosstalk/bleeding through. AT8 stained images in Cy5 were transformed to the red color, in place of magenta, to clearly show degree of co-localization. Triple fluorescent labelling was conducted using secondary antibody combinations without cross-reactivity, (anti-rabbit F(ab’)2-AlexaFluor-488, anti-rat F(ab’)2-AlexaFluor-555 and anti-mouse F(ab’)2-AlexaFluor-647) where selected visualization colors for imaging filters were applied as follows; FITC channel, green; TRITC channel, red; and Cy5 channel, magenta. For each case (tauopathy and control), the pertinent sub-pathologies for each disease were analyzed for their relationship with the candidate marker to provide comprehensive validation of cell-type (i.e. neuronal vs. astrocyte vs, oligodendrocyte) or cell-compartment specific aggregates (i.e. NTs vs NFTs, or APs vs. TAs). Co-localization is described as partial (i.e. decoration of distinct parts of aggregates) or full (majority/complete) co-localization. Enriched association in proximity to specific aggregates is as described. Tau neuropathological nomenclature in the current study refers to the following when describing results of proteomic candidate validation - by either proximal association, absence or degree of co-localization; NFT, neurofibrillary tangle; PT, pre-tangle; GLO/BN, globose/ballooned neuron in PSP; BN, ballooned neuron in CBD; NP, neuropil thread; DN, dystrophic neurites; NP, neuritic plaque; TANC, tangle associate neuritic cluster; AP, astrocytic plaque; TA, tufted astrocyte; CB, coiled body; GT, glial tau in PiD; VTs, VPS35 positive threads in PiD. Where possible, FFPE candidate validation was conducted in two or more cases for each disease, where each experiment included aged and young controls and secondary antibody only negative controls (see Supplementary Table 1 for case utility).

## Results

### ProPPr assay overview

For labelling and purification of AT8 phospho-tau-associated proteomes from post-mortem human tissue we utilized an antibody-driven horseradish peroxidase (HRP)-mediated proximity modification of proteins with biotin-tyramide molecules followed by isolation of modified proteins with streptavidin beads (Fig. 2A). Briefly, phospho-tau aggregates in 5 μm post-mortem FFPE brain tissue are immunostained with AT8 antibodies and HRP-conjugated anti-mouse secondary antibodies using standard immunohistochemical procedure. HRP is then used to catalyze the proximity modification of proteins that are adjacent to sites of antibody binding with biotin-tyramide. After stripping the antibodies, tissue is scraped out of the slides and lysed with partial formaldehyde cross-link reversal. Modified proteins are then isolated with streptavidin beads and analyzed by mass spectrometry.

To evaluate the efficiency and specificity of the ProPPr assay we used frontal cortex sections from 2 separate AD cases. In each of the cases we used 6 slides for labelling the phospho-tau-associated proteome as described above, while other 6 slides from the same case underwent the same procedure but without incubation with AT8 primary antibodies. These specific quality controls were conducted for each case to confirm enriched labelling in antibody positive samples versus antibody negative samples, in addition to and high signal to noise ratio from the pull-down of isolated proteins (Fig. 2B). Specificity of biotinylation was assessed by western blot analysis of tissue lysates with fluorescent streptavidin. We observed strong signal in the samples that were incubated with AT8 antibodies, but not in the samples where these antibodies were omitted, indicating that the labelling is dependent on the antibody staining and is thus specific. To evaluate the amount of protein isolated from these lysates by streptavidin beads, proteins were eluted from the beads, separated by SDS-PAGE followed by silver staining of the gel. We observed a drastic difference in protein amounts between samples that were subject to AT8 antibody incubation versus those with primary antibody omission, further indicating the efficiency and specificity of the ProPPr assay (Fig. 2B). Further quality controls include assessing the spatial distribution of NeutrAvidin-DyLight 488 labelling, which demonstrates the robust signal closely resembled that of the AT8-DAB IHC staining pattern of adjacent tissue slides from the same case (Fig 2B-F). The laminar distribution of AT8 pathology in AD cortex, in layers III and V of the neocortex is similarly highlighted by digital neuropathology color deconvolution quantification. Selection and stratification of cases were based on proof of principle studies and generation of the AT8 tau ProPPr cohort. For assay validation from all cases labelled, we performed *in situ* proximity labelling of phospho-tau-associated proteins in AD, CBD, PiD and PSP post-mortem frontal cortices and carefully assessed the specific neuropathology that was labeled by the ProPPr method (Fig 2C-F). We did not observe any significant signal in the cognitively unaffected control cases (Supplementary Fig. 1). Spatial patterns of biotin-tyramide modification were visualized by NeutrAvidin-DyLight 488 staining, and all disease-specific tau inclusions were fluorescent labelled, indicating high specificity and inclusivity of proximity labelling.

### Comprehensive profiling of phospho-tau-associated proteins by ProPPr

ProPPr was utilized to profile phospho-tau-associated proteomes in post-mortem frontal cortices across four major tauopathies. The association probability score of p-tau associated proteins was determined separately for each disease group using Mass spectrometry interaction Statistic (MiST) analysis, employing summarized protein intensity values from positive samples and their corresponding antibody-omitting negative controls. Using a MiST score cut-off value of 0.75 we identified 1318 phospho-tau-associated proteins, corresponding to 1315 unique protein-coding genes (Supplementary table 2). Predominantly, 1135 protein hits were identified as phospho-tau-associated proteins in AD cases, with 635 proteins in CBD, 699 proteins in PiD and 496 proteins in PSP cases (Fig. 3A, 3B).

**Fig. 3.**
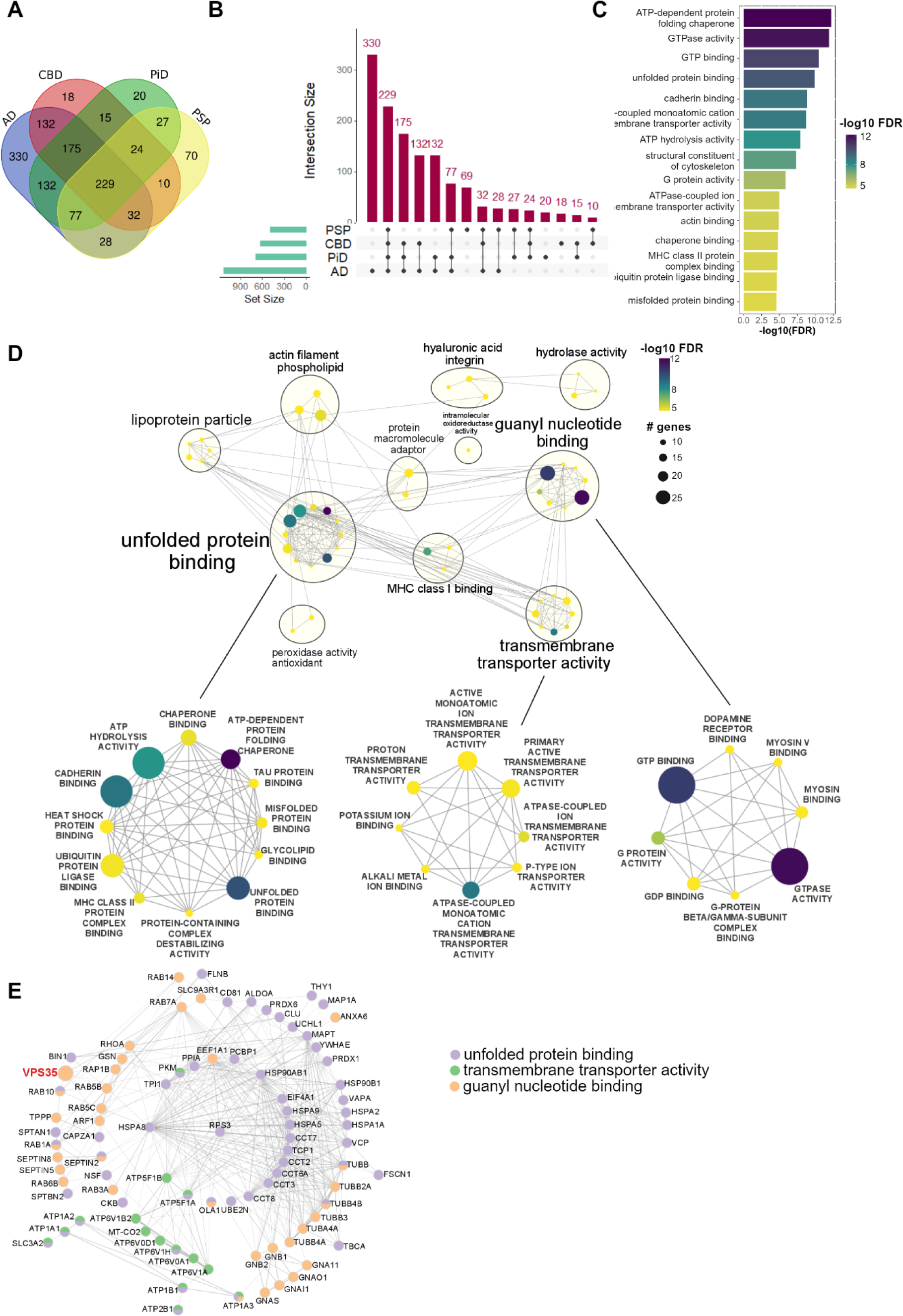
Overview of phospho-tau associated proteome across tauopathies. **(A**) Venn diagram for overlaps between the identified phospho-tau-associated proteins in tauopathies. **(B)** Upset plot for overlaps between the identified phospho-tau-associated proteins in tauopathies. **(C)** Top-15 GO terms enriched in the set of phospho-tau associated proteins identified in all the tauopathies (229 proteins). **(D)** Enrichment map network of top-50 GO terms enriched in the set of phospho-tau associated proteins identified in all the tauopathies (229 proteins). Clusters of nodes were annotated with the Autoannotate Cytoscape plugin. Color of the node represents false discovery rate (FDR), size of the node corresponds to the number of genes in GO terms. Lower panel contains close-up on the largest clusters of the enriched GO terms. **(E)** STRING network of protein-protein interactions for proteins from “unfolded protein binding”, “transmembrane transporter activity” and “guanyl nucleotide binding” clusters of GO terms.

To establish the concordance of our findings with established tau interactomes, we aligned and compared our p-tau associated protein dataset with previously reported tau interactors by utilizing a comprehensive review by Kavanagh *et al*., [73], which catalogues summarized information on 2084 identified physiological or pathological tau interacting proteins from 7 diverse experimental models and human brain studies (2035 unique gene names in total) [5, 41, 54, 61, 73, 104, 161, 167]. We observed a significant overlap, specifically that 844 of 1315 identified genes (∼64%) from the tau ProPPr dataset have been linked to tau interactome studies previously, showing statistically significant enrichment of known tau interactors in our results detailing phospho-tau-associated proteins (p-value =1.45*10^-568^ by hypergeometric test) (Supplementary Fig. 2A). Additionally, concordance analysis with a proteome of neurofibrillary tangles, isolated from AD patient brains by laser-capture microdissection [41], revealed 426 out of 542 proteins (∼78%) were also identified in our dataset, which comprises a statistically significant overlap (p-value =7.64*10^-421^ by hypergeometric test) (Supplementary Fig. 2B). Finally, we compared our protein sets with the recently published BAR-MS data on phospho-tau associated proteins in FFPE PSP tissue, despite differences in methodology [129]. This study reported 117 proteins (116 genes) associated with phospho-tau lesions in PSP, which is considerably lower than the 496 genes we report here, most likely due to the differences in the analysis pipeline and using more stringent criteria in the BAR-MS study. Nevertheless, our set of phospho-tau-associated proteins in PSP was significantly enriched in proteins identified in the PSP BAR-MS study (57 overlapped genes, p-value = 6.4 *10^-61^ by hypergeometric test) (Supplementary Fig. 2C). Our combined set of 1315 genes contained 102 out of 116 genes identified in the PSP BAR-MS study (p-value = 1.76*10^-106^ by hypergeometric test), suggesting that some of the phospho-tau-associated proteins identified in PSP brains may not be specific to PSP, but are rather common for many tauopathies (Supplementary Fig. 2D). Overall, this analysis affirms the reliability of our ProPPr assay in identifying known tau interactors and associated proteins, supporting its utility in profiling aggregate-associated proteomes in human tissues.

We proceeded to characterize the set of 229 common AT8 tau ProPPr associated proteins that we found consistently associated with phospho-tau across all studied tauopathies. Gene ontology (GO) term overrepresentation analysis highlighted “ATP-dependent protein folding chaperone” as the most significantly enriched term in this protein set, aligning with known involvement of chaperones in misfolded tau aggregates (Fig. 3C, Supplementary table 3). Visualization of the top-50 enriched GO terms as an enrichment map network revealed several clusters of closely related terms (Fig. 3D). The largest cluster of terms annotated as “unfolded protein binding” contained the above-mentioned top-overrepresented GO term and 11 additional clusters that included “unfolded protein binding”, “ATP hydrolysis activity”, “ubiquitin protein ligase binding”, “chaperone binding”, “heat shock protein binding”, “misfolded protein binding”, “tau protein binding”, “protein containing complex destabilizing activity terms”, “cadherin binding”, “glycolipid binding” and “MHC class II protein complex binding” (Fig. 3D). Identified phospho-tau-associated proteins that belong to this term include, among others, several heat shock proteins of HSP70 (HSPA1A/HSPA1B, HSPA2, HSPA5, HSPA8) and HSP90 families (HSP90AB1), chaperone-like proteins CLU and VCP, as well as constituents of CCT/TriC chaperonin complex (TCP1, CCT2, CCT3, CCT6A, CCT7, CCT8) (Fig. 3E).

An additional prominent cluster was annotated as “transmembrane transporter activity”, encompassing 8 GO terms primarily related to membrane ion transporters (Fig. 3G). Proteins from these terms included several subunits of vacuolar-type proton ATPase (ATP6V0A1, ATP6V0D1, ATP6V1A, ATP6V1B2, ATP6V1H), Na^+^/K^+^ ATPase subunits (ATP1A1, ATP1A2, ATP1A3), plasma membrane Ca^2+^-transporting ATPase ATP2B and mitochondrial ATP synthase subunits (ATP5F1A, ATPF1B) (Fig. 3E).

The remaining major cluster was annotated as “guanyl nucleotide binding” and contained 8 GO terms, notably including the terms “GTP binding” and “GTPase activity” as the largest and the most significantly overrepresented terms in the analysis (Fig. 3D). Proteins from these terms included several members of the RAB GTPase family (RAB1A, RAB3A, RAB5B, RAB5C, RAB6B, RAB7A, RAB10 and RAB14), G protein subunits (GNAI1, GNA11, GNAO1, GNAS, GNB1, GNB2), septin proteins (SEPTIN2, SEPTIN5, SEPTIN8), and several tubulin proteins (TUBA4A, TUBB, TUBB2A, TUBB3, TUBB4A, TUBB4B) (Fig. 3E).

### VPS35 is sequestered into heterogenous phospho tau aggregates across tauopathies

Among the p-tau associated proteins shared across tauopathies and falling within the “guanyl nucleotide binding” cluster we identified VPS35 (vacuolar protein sorting-associated protein 35) (Fig. 3E). VPS35 is a member of a cargo-selective complex with VPS26 and VPS29 and constitutes an integral part of the retromer complex crucial for endosomal trafficking, facilitating transport of selected cargos from endosomes to the trans-Golgi network or plasma membrane [141]. The retromer complex, including VPS35 has been implicated in several neurodegenerative diseases [169]. Most notably, the D620N mutation in *VPS35* gene is causative of late-onset Parkinson’s disease in an autosomal-dominant manner [180]. Given these strong links, we decided to explore the association of VPS35 with phospho-tau lesions by co-immunostaining post-mortem frontal cortex sections from tauopathy patients with AT8 and VPS35 antibodies (Fig. 4).

**Fig. 4.**
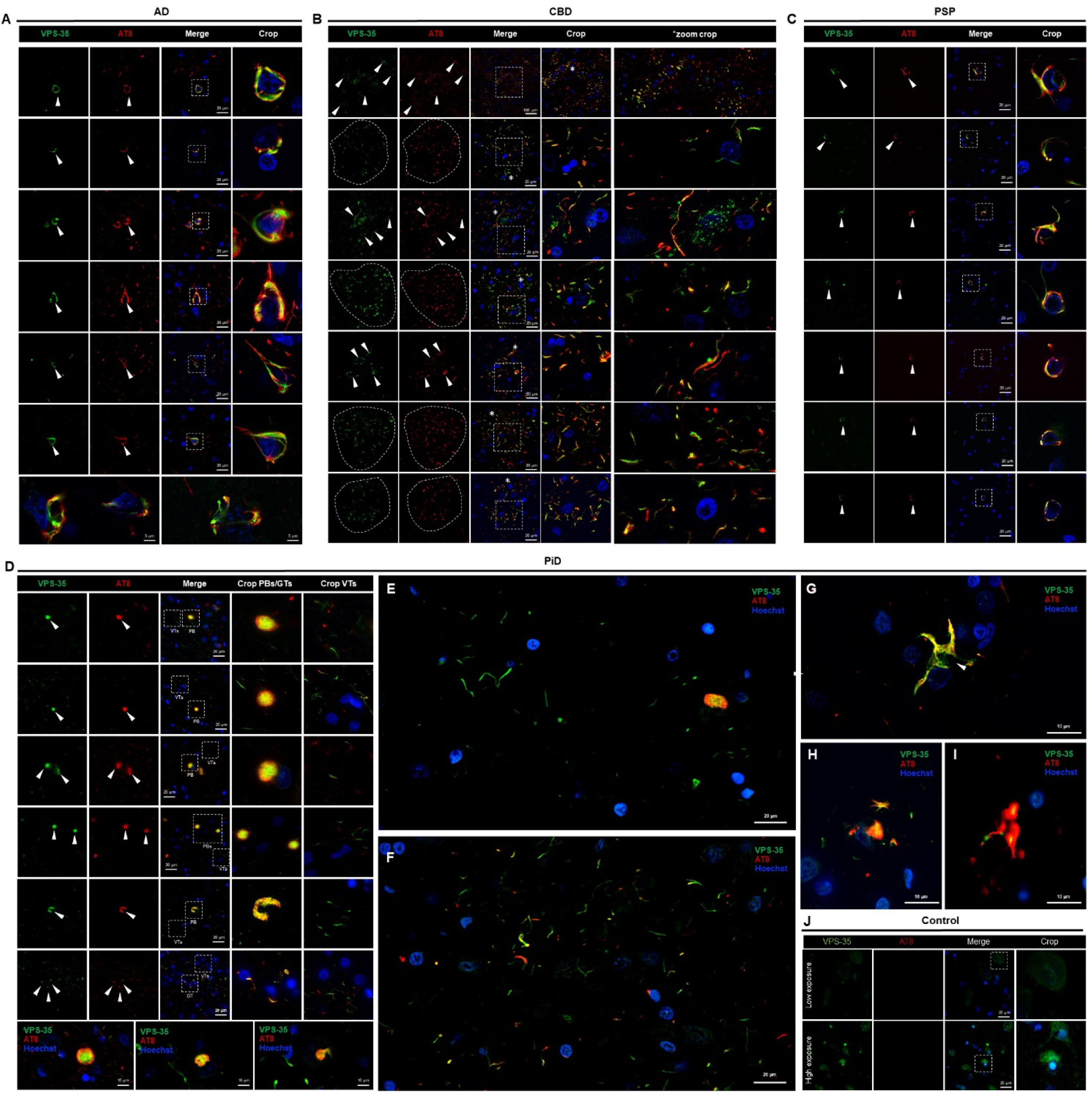
VPS35 co-localizes with specific disease related AT8 tau-positive pathologies in Alzheimer’s disease and primary tauopathies. Post-mortem frontal cortex (BA9) from tauopathy and control cases were labelled with VPS35 (green) and AT8 (phospho-tau-red) via immunofluorescence, which depict distinct pathologic tau aggregates that co-localize with abnormal VPS35 accumulation amongst AD and primary tauopathy cases. Images show nuclear staining with Hoechst (blue) and merged and cropped images as labelled. **(A)** Neurofibrillary tangles and pre-tangles from Alzheimer’s disease cases reveal robust co-localized immunostaining with VPS35. Neuropil threads and neuritic plaques are absent for VPS35 colocalization. **(B)** Hallmark pathologies of CBD, astrocytic plaques, show co-localized signal of VPS35 and AT8 from early to advanced sized plaques, as indicated by arrows and circular dashed regions of plaques. Thick neuropil thread pathology also shows corresponding signal between both markers. **(C)** Oligodendroglial inclusions in PSP demonstrates that AT8 positive coiled bodies are co-positive for VPS35, yet tufted astrocytes and globose NFTs do not form the same pathologic association. **(D)** Circumscribed neuronal Pick bodies (PBs) display VPS35 punctate staining within the core at AT8 positive inclusions and a distinct phenomenon of frequent VPS35 threads (VTs) negative for AT8 tau **(E)**, which were occasionally partially co-localized with AT8 immunostaining **(F)**. Glial tau in Pick’s disease **(G-I)** displayed full to partial co-localization of AT8 and VPS35. **(J)** Unaffected control tissue denoting two exposures of AF-488 channel, low exposure akin to disease tissue sections (100 msec) or over exposure (700 msec) denoting the non-disease distribution of VPS35 in a cytosolic vesicular form. Abbreviations: AD, Alzheimer’s disease; AT8, phospho-tau pS202/Thr 205; CBD, corticobasal degeneration; PSP, progressive supranuclear palsy; PiD, Pick’s disease; CTRL, control; VPS35, vacuolar protein sorting ortholog 35; *, zoom crop. Scale bars denoted in each figure pane.

In AD frontal cortices, VPS35 demonstrated robust co-localization with AT8 positive pre-tangles and mature neurofibrillary tangles (Fig. 4A). Conversely, other phospho-tau lesions typical of AD pathology, such as neuritic plaques and neuropil threads were mostly devoid of VPS35 signal. In CBD frontal cortex, VPS35 exhibited strong co-localization with astrocytic plaque pathology and some larger neuropil threads (Fig. 4B). In PSP frontal cortices, all oligodendroglial coiled bodies were positive for VPS35 signal (Fig. 4C). However, tufted astrocytes and globose neurofibrillary tangles did not display co-localization with VPS35. In PiD, VPS35 was largely sequestered into intraneuronal Pick bodies, displaying punctate staining within their cores (Fig. 4B). Additionally, VPS35 signal was evident in smaller AT8 lesions, likely representing rarer glial phospho-tau aggregates (Fig. 4D). Intriguingly, in PiD we observed frequent thread-like VPS35 structures that were often negative for AT8 phospho-tau (Fig. 4E) or only partially co-localized (Fig. 4F), a phenomenon not observed in cognitively unaffected individuals or patients with other tauopathies. Glial tau pathology ranged from full to partial co-localization (Fig. 4G-I), whereas control tissue displayed normal VPS35 staining at far lower intensity than VPS35 sequestered into AT8 lesions. The precise nature of these predominantly VPS35 positive threads and their potential specificity to PiD warrant further investigation.

Validation of candidate proteins by high-resolution co-immunofluorescence of post-mortem frontal cortices corroborates the mass spectrometry proteomics finding, confirming VPS35 association with AT8-positive aggregates in all four major tauopathies. Nonetheless, we also demonstrate the selective extent of VPS35 sequestration among different phospho-tau lesions in distinct diseases.

### ProPPr identifies differential association of proteins with phospho-tau aggregates between tauopathies

The MiST algorithm implemented in an online web tool, was used for the identification of phospho-tau associated proteins. However, since it provides a “yes/no” type of answer to whether particular proteins interact with a studied bait based on the cut-off applied, it is not designed for quantitative comparisons of protein-protein interactions between several conditions. To find proteins with altered association with phospho-tau lesions between AD, CBD, PiD, and PSP, we performed statistical comparison of protein abundances in samples from corresponding cases using DEP R package. We identified 31 proteins with significant change in abundance (adjusted p value <=0.05) in at least 1 of 6 pairwise comparisons of studied disease conditions. Clustering analysis based on Pearson correlation revealed that the samples from cases with the same pathological diagnosis generally clustered together, indicating that abundances of the identified significantly changed proteins can be characteristic features that help distinguishing different types of tau pathology (Fig. 5A). Significantly changed proteins are distributed along the range of protein abundances inferred from mass spectrometry data, suggesting that our analysis was not biased towards highly abundant or low abundant proteins (Fig. 5C). Most of the changes were detected in comparison between PiD and PSP (19 differentially abundant proteins), followed by comparison between AD and CBD (11 proteins), AD and PSP (4 proteins) and CBD vs PiD (1 protein). Comparisons between AD and PiD, as well as CBD vs PSP did not reveal any statistically significant protein differences in our data (Fig. 5B).

**Fig. 5.**
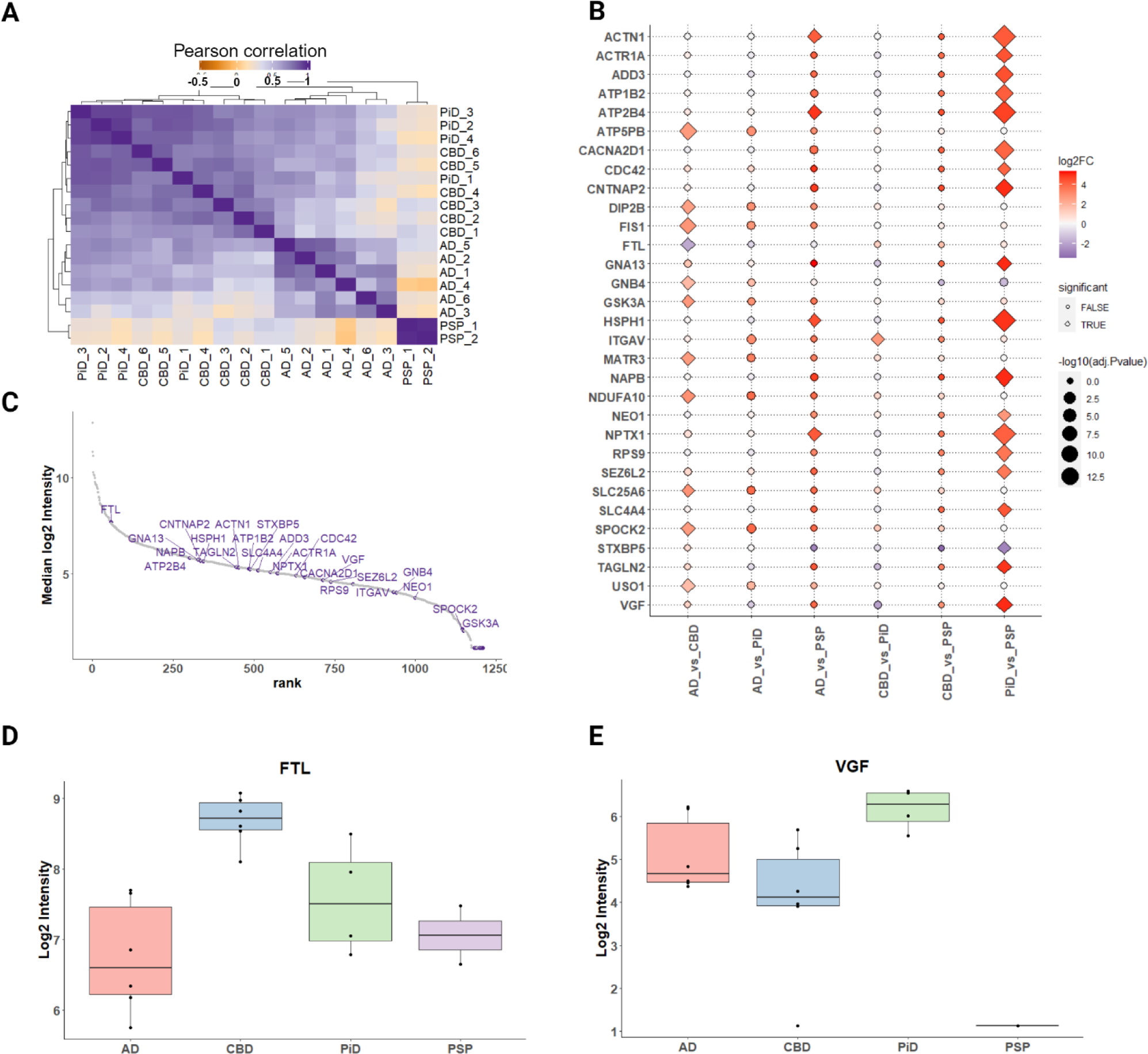
Analysis of differential protein association with phospho-tau lesions across tauopathies. **(A)** Sample correlation plot generated using abundance values for the proteins significantly changed in at least one of the pairwise comparisons between the tauopathies. **(B)** Log2 fold changes for proteins with significant changes in association with phospho-tau in each pairwise comparison presented as a dot plot. Diamond shape indicates statistically significant difference (adjusted p-value <0.05). **(C)** Dynamic range of median protein intensities in the mass spectrometry proteomics data. Significantly changed proteins are shown in purple. **(D)** Summarized FTL protein intensity in tauopathies measured by mass spectrometry. **(E)** Summarized VGF protein intensity in tauopathies measured by mass spectrometry.

No significantly overrepresented pathways or GO terms were identified among the set of proteins with differential abundance between tauopathies. Consequently, we focused on individual proteins previously linked to neurodegenerative diseases. Particularly, neurosecretory protein VGF has emerged in large-scale omics studies as a potential biomarker of several neurodegenerative and psychiatric diseases [128]. We observed a significant increase in VGF levels in neuropathologic aggregates of PiD compared to PSP, with a specific trend for pathologic VGF enrichment in PiD to be compared to other tauopathies (Fig.5B, 5E). Another notable protein was ferritin light chain (FTL). Mutations in the *FTL* gene lead to neuroferritinopathy, an autosomal dominant neurodegeneration within brain iron accumulation disorder [28, 97, 165]. FTL was highlighted as the highest abundant protein amongst those analyzed in our dataset and was found to be majorly sequestered into CBD pathology, with a general trend to be elevated in CBD compared to AD (Fig. 5B, 5C, 5D). These findings suggest that discernible differences in phospho-tau-associated proteomes between tauopathies can be detected by mass spectrometry proteomics and warrant careful neuropathologic analysis of disease-specific tau sub-pathology to identify the origin and selectivity of the enrichment.

### VGF is associated with Pick bodies and Alzheimer’s disease neurofibrillary pathology

In our proteomic analysis, the neurosecretory protein VGF exhibited a preferential association with phospho-tau lesions in PiD. To validate this finding, we performed co-immunofluorescent imaging of VGF with phospho-tau in tissue from the four major tauopathies (with AD, CBD, PiD, and PSP) and cognitively unaffected individuals (Fig. 6). In AD cases the number and density of VGF puncta in the neuropil were visibly reduced. In certain small NFTs, VGF decorated AT8 positive inclusions (Fig. 6 B-C), whereas in pre-tangles (Fig. 6D) and TANCs (Fig. 6E), VGF signal partially co-localized, suggesting active association of VGF and sequestration into these pathologic tau structures. Of note, we also observed proximal and interspersed enrichment of VGF staining in the environment of AD neuritic plaques (Fig. 6A). Although not directly co-localized with AT8-positive neurites, this hallmark requires further investigation to delineate the relationship with the amyloid component of these types of AD plaques, specifically owing to the dual proteopathic nature of these lesions. This result also suggests that in addition to NFTs, VGF may mediate specific accrual relationships within mature amyloid-loaded AD neuritic plaques, as has been previously reported [40]. There was no association with VGF and AD neuropil threads, indicating that this pattern is likely associated with the maturation level and cellular compartment specific pathologies in the context of AD pathogenesis.

**Fig. 6.**
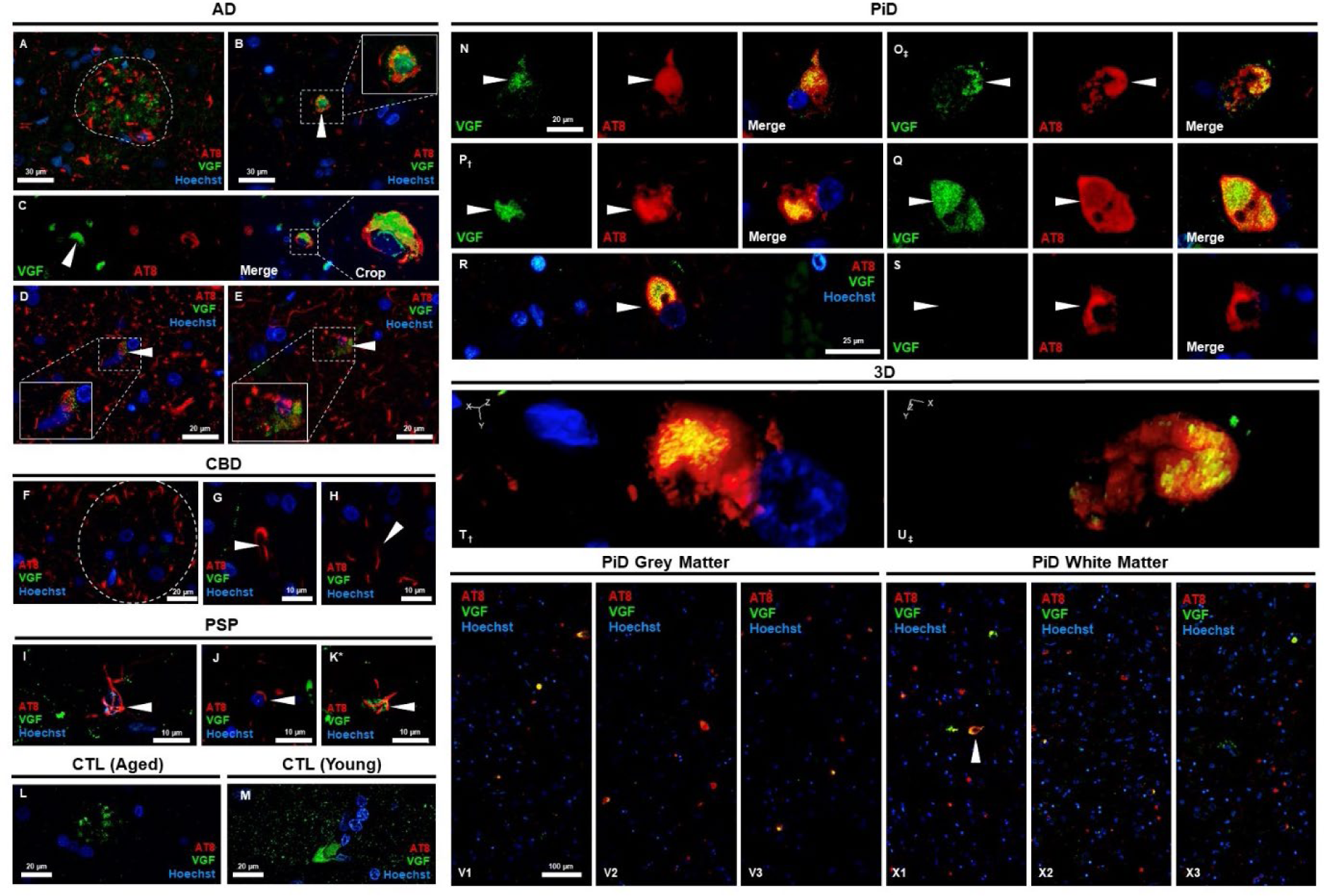
VGF colocalizes with distinct pathological tau aggregates in the frontal cortex of human post-mortem brain tissue. Co-immunofluorescent staining with VGF (green) and AT8 phospho-tau (red) was conducted on FFPE human brain tissue from Alzheimer’s disease, primary tauopathy and control cases, which were absent for tau pathology in the frontal cortex. VGF was enriched within tau positive neuritic plaques **(A)** and co-existed as punctate signal decorating small neurofibrillary tangles and pretangles **(B-D)** and tangle associated clusters **(E)**. VGF did not co-localize in astrocytic plaques of CBD **(F)**, oligodendroglial coiled bodies **(G)** or neuropil thread pathology **(H)**. Most PSP tufted astrocytes and coiled bodies did not co-localize with VGF signal **(I,J)** and rare partial co-localization signal (yellow) was observed in a minority of tufted astrocytes **(K*)**. The distribution of VGF observed in control tissue from age-matched **(L)** and young **(M)** cases, which both lacked phospho-tau pathology in frontal cortex, revealed a punctate signal enriched highly within some cells. Both spherical, coned and horseshoe shaped Pick bodies in PiD cases demonstrated the highest and most robust co-localization of AT8 tau and VGF **(N-R)** . Further, PiD glial inclusions sporadically observed in PiD cases, were either not co-positive **(S)** or positively co-localized in white matter **(X1)**. 3D projection images of rendered z-stacks images revealed the depth of VGF co-localization from pick bodies in **(T, U)**, which is usually at the core of pick bodies. 60x tile images revealed the distribution and degree of VGF positive pick inclusions in the grey **(V1-V3)** and white matter **(X1-X3)**. Scale bars denoted in figures. Abbreviations: AD, Alzheimer’s disease; AT8, phospho-tau pS202/Thr 205; Scale bars denoted in each figure pane; CBD, corticobasal degeneration; PSP, progressive supranuclear palsy; PiD, Pick’s disease; CTRL, control; *, rare phenomena; †, full focused z-stack and 3D rendering pair (V and T); ‡ full focused z-stack and 3D rendering pair (O and U).

We did not find any significant co-localization between VGF and CBD astrocytic plaques, coiled bodies, globose NFTs or threads (Fig. 6F-H) or in the majority of PSP tufted astrocytes (Fig.6I), coiled bodies (Fig. 6J), or neuronal pathology. There were rare occasions where small tufted astrocytes were partially co-localized with VGF puncta (Fig. 6K). In PiD we also observed a reduction in the number of VGF puncta present in the cellular milieu of the neuropil compared to control tissue. Strikingly, we found strong VGF signal in the majority of the spherical, coned and horseshoe-shaped Pick bodies found in PiD frontal cortex grey matter (Fig.6 N-R,V1-V3). In specific cases where Pick pathology was severe and extended through to the white matter, we did not observe such a strong association with VGF accumulation (Fig. 6X1-X3). VGF was located in the center core of Pick bodies and encased by AT8-positive tau. Upon 3D rendering of select PiD lesions, we saw that the co-localized signal of AT8 and VGF was indeed centralized and encapsulated by tau, even within 5 µm tissue sections (Fig.6 T,U). Rarer glial inclusions in the grey matter lacked co-localization with VGF (Fig. 6S), while some white matter glial inclusions showed co-localization (Fig.6 X1, arrow). Thus, our immunohistochemical analysis corroborates the proteomics data, indicating a predominant VGF association with pathologic tau in PiD. Control tissue revealed a punctate distribution of VGF in the neuropil, with occasional cells showing strong VGF signal in soma (Fig.6 L-M), interestingly young control tissue (Supplementary Table 1. Case #35) had high levels of VGF distributed amongst the grey matter over aged cases.

### FTL-positive microglia are associated with phospho-tau in CBD astrocytic plaques

The results of our proteomic data analysis point to a strong association of FTL with specific tauopathies and their defining sub-pathologies. In AD enrichment of FTL was observed, despite lack of co-localization with AT8 pathology. Interestingly, AD AT8 structures, including NFTs, PTs, NPs and NTs were not co-positive but were outweighed by the high intensity FTL accumulation within small cellular structures resembling microglia (Fig. 7). Accompaniment of highly positive FTL somal compartments of cells and within neuritic plaques, were also identified in AD. Yet the degree of signal localizing on small, recruited cells adjacent to AD pathology aligns with the known disease-associated microglial involvement of AD [55]. In PiD, FTL was not detected in Pick bodies but engaged within small phospho-tau-positive structures that most likely represent low frequency and 4R-tau positive glial tau inclusions [21, 80, 84]. FTL could also be detected in PSP tufted astrocytes, although its co-localization with these lesions was not as striking as with astrocytic plaques in CBD. Coiled bodies showed co-localization whereas no FTL was present in PSP globose NFTs.

**Fig. 7.**
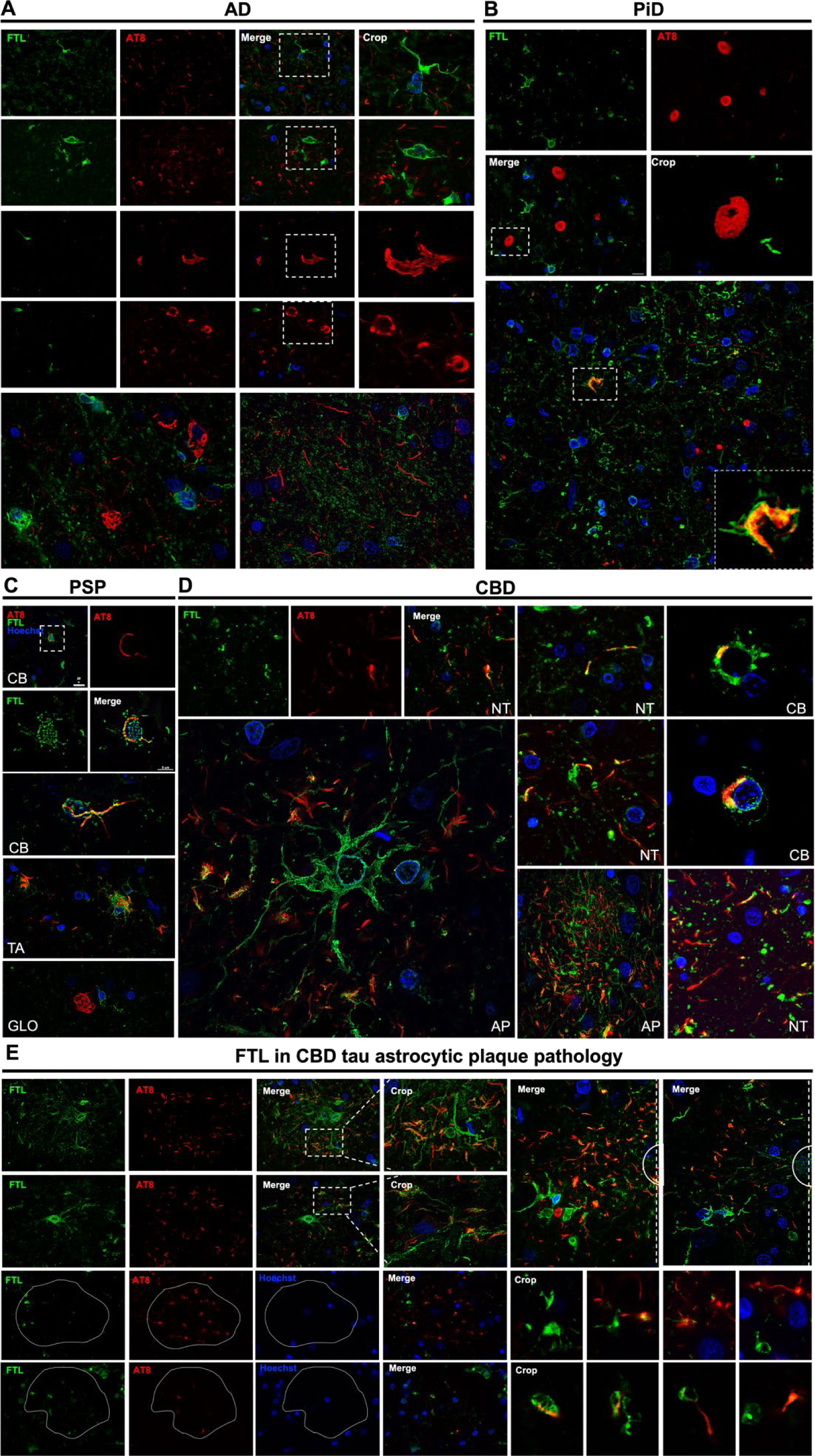
FTL colocalizes with glial tau pathology in CBD, PSP, and Pick’s disease. Co-immunofluorescent staining with FTL (green) and AT8 phospho-tau (red) was conducted on FFPE human tauopathy brain tissue. **(A)** In AD, FTL was not associated with NFT, PT or NT pathology, but was enriched in microglial cells associated with neuritic plaques. **(B)** Pick bodies were not-colocalized with FTL signal, but rare glial phospho-tau inclusions in the white matter were positive for FTL. **(C)** PSP coiled bodies and tufted astrocytes co-localized with FTL signal, as demonstrated by decoration of FTL puncta on coiled bodies and within the projections of AT8 tufted astrocyte inclusions. Globose NFTs in PSP cases were not stained with FTL. **(D)** The distribution of FTL in lesions of CBD frontal cortex tissue demonstrated strong co-localization in the distal extensions of phospho-tau APs and coiled bodies, with partial co-localization demonstrated in AT8 positive neuritic threads. **(E)** Examination of CBD astrocytic plaque pathology revealed multiple cellular components that were FTL positive and differentially associated with small and larger APs. Larger and more mature APs (upper left panels) showed extensive and dystrophic FTL positive cells in the center core and recruitment to the outer perimeter (upper right panels), with semicircles denoting the core region of the APs. Smaller and less mature APs (lower left panels) displayed smaller and more rounded morphology of recruited FTL positive cells that were also partially co-localized with phospho-tau (lower left panels). Abbreviations: AD, Alzheimer’s disease; AT8, phospho-tau pS202/Thr 205; CBD, corticobasal degeneration; PSP, progressive supranuclear palsy; PiD, Pick’s disease; NFT, neurofibrillary tangle; NP, neuritic plaque; CB, coiled body; TA, tufted astrocyte; AP, astrocytic plaque; GLO, globose NFT; NT, neuropil thread.

In CBD cases compared to AD and other tauopathies, co-immunofluorescence of frontal cortex with AT8 and FTL revealed full co-localization of FTL with distal extensions of astrocytic plaques, in addition to neuropil threads, and coiled bodies (Fig. 7D-E). Upon examination of astrocytic plaques of different sizes, likely reflecting the distal progression of tau accumulation within these hallmarks, we noticed that smaller astrocytic plaques often don’t demonstrate robust co-localization with FTL that is more characteristic of more mature plaques. However, in most cases we could observe the presence of FTL-positive cells in the vicinity of these small astrocytic plaques (Fig.7E). This may suggest that FTL co-localization with astrocytic plaques may not result from FTL upregulation only in astrocytes affected by tau aggregation, but from FTL delivery to forming astrocytic plaques by other cells that respond to tau pathology, such as microglia.

To test this suggestion, we performed triple co-immunofluorescent labelling of CBD frontal cortices with AT8, anti-FTL antibodies, and antibodies to either astrocyte (GFAP) or microglia (IBA1) cell-type specific markers. Surprisingly, we found that AT8 signal co-localized with FTL in large and mature astrocytic plaques, which were also co-positive for IBA1 (Fig. 8A-D), whereas in some smaller lesions, GFAP co-localization was mainly observed, albeit with FTL positive cells in the vicinity (8E-G). IBA1 and FTL positive microglia demonstrated strong co-localization and center cells highlighted the extensive level of dystrophic morphology. In a wide view image of multiple CBD lesions found in the frontal cortex (Fig. 8D), IBA1/FTL co-positive cells dominated larger astrocytic plaques and also show active and motile phenotypes closely associated to regions of phospho-tau pathology. Astrocyte marker analysis of these larger APs (Fig. 8F) reveled fragmented GFAP positive extensions, lacking an observable somal compartment, and instead show an FTL positive cell in the center core. This data indicates that plaques may somehow recruit microglia and induce transformation to an FTL-positive microglial phenotype that colonizes the astrocytic plaque during the disease progression of CBD and maturation of pathology.

**Fig. 8.**
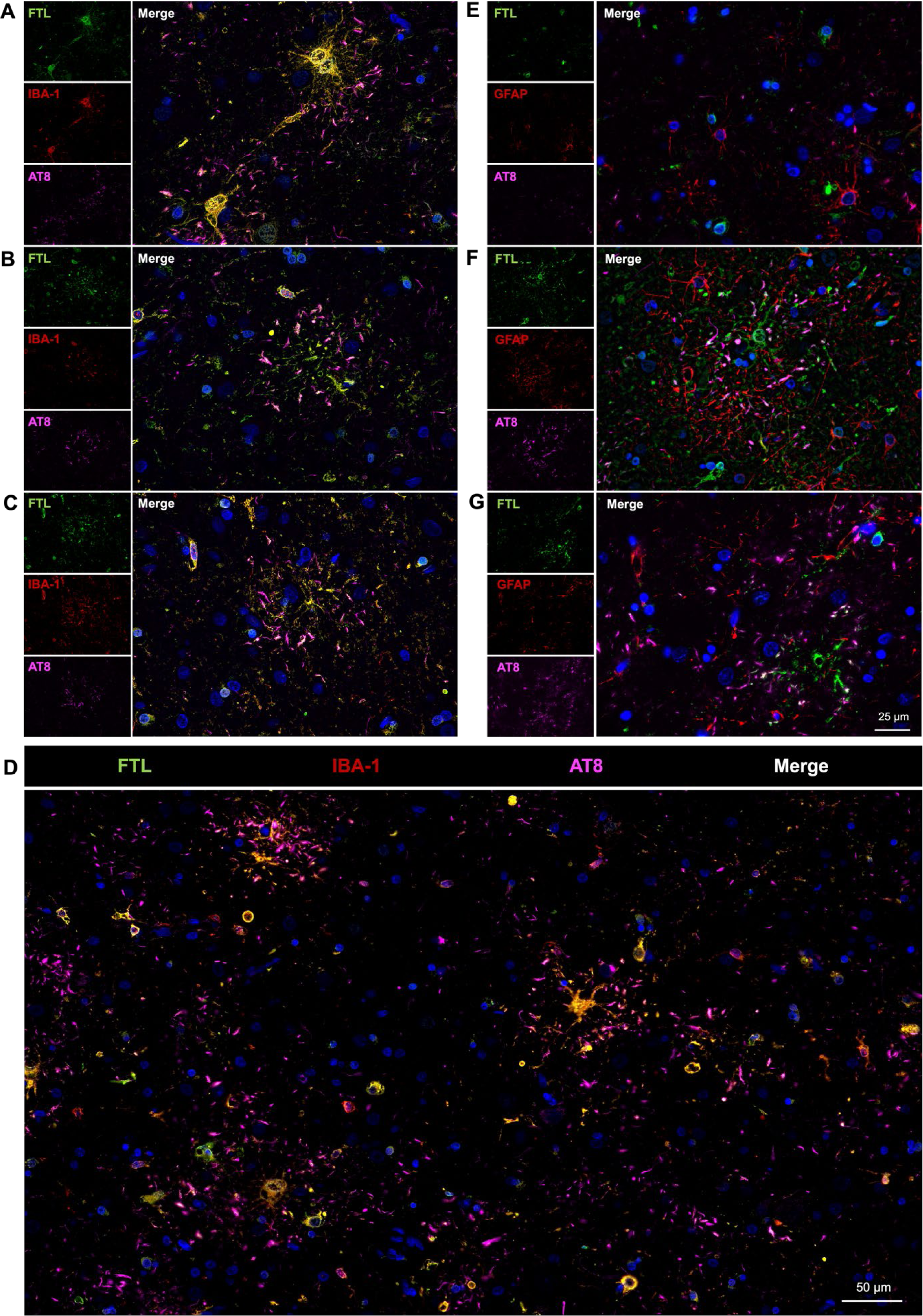
CBD astrocytic plaque triple co-immunofluorescence with glial markers show the microglial component of these tau lesions in association with increased expression of FTL. Closer examination of the glial components of CBD AP tau pathology with IBA1 or GFAP (red), FTL (green) and AT8 phospho-tau (magenta), highlighted the enriched involvement of microglia in CBD astrocytic plaques. (**A-C**) FTL was strongly co-localized with IBA1 positive microglia that were recruited to these aggregates. In larger plaques IBA1 positive cells were localized to the central core dominating the area of the principal cells emanating from APs. The highly dystrophic and thickened morphology of FTL positive microglia revealed close association with the APs and co-localization with the distal extensions of the tau positive corona of APs. (**D**) High resolution tile imaging reveals the distribution of FTL and astrocytic plaques, including both dystrophic and amoeboid FTL microglia. **(E)** Co-staining of FTL, GFAP and AT8 demonstrated GFAP positive astrocytes in very early states of CBD AP formation as shown by smaller and less mature AP densities. **(F-G)** In larger APs, GFAP positive extensions were present, but not a central somal component, whereas GFAP negative cells, positive for FTL were located closer to the core. **(G)** While some co-localization between GFAP and AT8 tau occurs in APs, the location of dystrophic FTL positive cells dominate this advancing maturation of pathology in CBD. Abbreviations: AT8, phospho-tau pS202/Thr 205; CBD, corticobasal degeneration; AP, astrocytic plaque.

## Discussion

Studies of interactome or associated proteome of pathological tau species provide important insight into mechanisms of tau-mediated toxicity in neurodegenerative conditions. In recent years, a number of research studies evaluated the interactome of tau in various cellular models using immunoprecipitation or proximity biotinylation approaches [5, 41, 54, 61, 68, 91, 99, 104, 149, 161, 166, 167]. These studies provided a comprehensive knowledge of tau interactors in normal and pathologic conditions that may function as active modifiers of tau by disrupting or inciting its aggregation, which leads to neurodegeneration. The common feature of co-immunoprecipitation studies is that the content of isolated protein complexes is highly dependent on buffer conditions used in these assays. On the other hand, the commonly used proximity proteomics methods such as BioID or APEX2 rely on expression of genetic constructs in cellular, tissue or animal models. While there are many models of tauopathy described in the literature, they do not fully recapitulate the heterogeneity of pathological tau lesions observed in human diseases, making the post-mortem disease human brain the only reliable source for studying tau aggregates in situ with a true disease specific nature. Methods such as laser-capture microdissection have provided invaluable information on the proteome of tau aggregates in disease but are only applicable for the dissection of large-sized lesions that are well defined spatially, such as the soma of NFTs or the circumscribed Pick bodies. Thus, development of methods that provide a capability to capture interactome or associated proteome of pathological protein aggregates such as the ones formed by tau protein *in situ* is critically required for understanding of molecular mechanisms of neurodegenerative pathology. In support of this, a recent study that used *in situ* proximity proteomics strategy for identification of phospho-tau-associated proteome in PSP cases identified 19 novel proteins that were not previously found to be associated with Tau [129]. Here we describe our ProPPr technique that utilizes a similar strategy but alternative protocol for proximity proteomics, and potentially other omics profiling. Since neuronal and/or glial tau aggregates are a common feature of tauopathies, it remains unknown how these inclusions emerge in specific cell types or compartments and develop across regionally vulnerable brain areas. The diversity of neuropathology supports the hypothesis that differential tau-associated proteomes will be discovered using our spatially conserved proximity labelling method, ProPPr, across major tauopathy diseases.

Using this method, we have performed the first comprehensive ProPPr profiling of the phospho-tau-associated proteome across four major tauopathies – AD, CBD, PiD, and PSP, in the same study and with relevant validation. While this strategy is related to various assays described before [7, 75, 129, 130], it has several important modifications. Firstly, instead of thick free-floating sections, for each studied case we used several thin 5 μm tissue sections mounted on standard glass slides, which are more easily available from brain banks, allow for automated workflows with autostainers, and are expected to increase the level of antibody penetration and protein labelling density. Secondly, for the tissue lysis and reversal of crosslinking we used buffers based on the MS-compatible surfactant RapiGest instead of commonly used SDS-containing buffers [119]. Finally, the utilization of biotin-SS-tyramide that contains disulfide bond cleavable by reducing agent allowed us to elute purified tyramide-modified proteins from streptavidin beads, potentially increasing the specificity of the assay.

We report identification of 1318 phospho-tau-associated proteins from 1315 genes, 471 of which were not reported in previous mass spectrometry proteomic studies investigating the tau interactome. While our study represents the first attempt of comparative analysis of phospho-tau-associated proteome between tauopathies, it is subject to certain limitations that require consideration. A modest sample size of cases in our experimental cohort likely led to an underestimation of the number of proteins exhibiting significantly different associations with phospho-tau aggregates among the tauopathies. Subsequent investigations involving larger cohorts may yield additional insights into both common and tauopathy-specific proteomes of pathologic tau lesions. Furthermore, advancements in the efficiency of the ProPPr assay can likely be further improved by utilization of newly introduced ultra-sensitive mass spectrometry instruments for single-cell proteomics and refined techniques of protein extraction from FFPE tissue, such as focused ultrasonication [127]. Nevertheless, our study provided a comprehensive overview of phospho-tau-associated proteome in tauopathies and uncovered several important findings that provide novel insight towards the pathological events and features that are common or specific to major tauopathy diseases examined here.

Among the identified 1318 phospho-tau-associated proteins 229 proteins are associated with phospho-tau in all four major tauopathies: AD, CBD, PiD, and PSP. The largest group of interrelated GO terms overrepresented in this 229-protein list is associated with unfolded protein binding, including the most significant “ATP-dependent protein folding chaperone” term. These terms include proteins that belong to several groups of molecular chaperones: HSP70, HSP90, CCT/TriC, and chaperone-like proteins CLU and VCP. The proteins from these groups, including the ones identified by us, were previously found in several studies of tau interactome and neurofibrillary tangle proteome [41, 73]. Moreover, CLU has been previously shown to co-localize with tau deposits in AD, CBD, and PiD post-mortem brains, and HSP70 and HSP90 chaperones were found to be sequestered in seeding-induced tau aggregates *in vitro* [170, 173]. Recruitment of molecular chaperones into phospho-tau aggregates likely reflects the response of cellular folding machinery to emergence of misfolded tau species and its attempts to re-fold tau molecules or send them for degradation. It has been shown that different chaperones can effectively reduce tau phosphorylation, filament formation and solubility [33, 38, 123, 124, 170]. However, it has also been demonstrated that chaperone-activity can stabilize amyloidogenic tau folding, facilitate tau filament formation or increase production of seeding-competent tau forms, promoting spread of tau pathology [12, 27, 69, 114, 136, 160, 176]. Thus, association of molecular chaperones with tau lesion may have both positive and negative effects on aggregation of tau and progression of tau pathology. In any case, sequestration of molecular chaperones in tau aggregates likely aggravates proteostasis imbalance.

Other notable groups of phospho-tau-associated proteins include the subunits of V-type ATPase, RAB GTPase family members, G protein subunits and Septins. Many of them have been found in tau deposits in human brain previously, but the functional consequence of their sequestration in tau aggregates has not been studied extensively [41, 77, 171]. Additionally, little is known about potential roles of these proteins in tau pathogenesis, except for the role of RAB7A in tau secretion [132]. Functional relationships between these proteins and tau aggregation may be exciting themes for future studies. For example, it is tempting to hypothesize that sequestration of the subunits of the V-type ATPase in tau deposits disrupts acidification of endosomes and lysosomes, leading to defects in endolysosomal system – a well-known phenomenon in AD and other neurodegenerative diseases [125, 178].

One of the proteins found associated with tau in all four major tauopathies is core retromer complex protein VPS35. This protein has been previously found in the proteome of neurofibrillary tangles in AD and identified as tau interactor in several studies of tau interactome [41, 61, 68, 161, 166]. We validated the association of retromer complex protein VPS35 with several types of phospho-tau lesions in frontal cortices from all the studied tauopathies by fluorescent immunohistochemistry. Although the acute effect of tau aggregation on retromer complex activity has not yet been directly evaluated, it seems plausible that sequestration of VPS35 in insoluble tau aggregates may lead to retromer complex deficiency. In support of this, several lines of evidence suggest the impairment of retromer functioning in tauopathies. Retromer dysfunction in AD is indicated by reduction of VPS35 and VPS26 protein level in AD patients’ entorhinal cortex that is particularly vulnerable for tau pathology in AD [150]. Similar phenomenon was also observed in brains of young and aged Down syndrome patients that have increased risk of developing AD neuropathology after age of 40 [29]. In addition, the known retromer complex receptor SORL1 is decreased in trans-entorhinal cortex in AD, which may be caused by insufficiency of SORL1 recycling by retromer [148]. Finally, retromer complex is known to play an important role in regulating lysosomal homeostasis [30], and lysosomal defects are well documented phenomena in both AD and primary tauopathies, although the exact mechanisms that lead to this effect are not entirely clear [125, 178]. Altogether, these data suggest that retromer complex disfunction is a characteristic feature of tauopathies, and VPS35 sequestration in phospho-tau pathological lesions may be one of the mechanisms that underlies this effect. Importantly, there is numerous evidence that shows detrimental effect of retromer dysfunction and beneficial effect of restoration of the retromer complex function on tau pathology [17, 18, 20, 51, 87, 172]. This may suggest the existence of positive feedback loop between tau aggregation and retromer disfunction.

Development of the ProPPr assay allowed us to perform a first comparative analysis of phospho-tau-associated proteome between four major tauopathies. Selected candidates with differential association with phospho-tau have been validated by immunofluorescent staining of post-mortem patient frontal cortices followed by microscopic examination of stained tissue. Using this approach, we demonstrated sequestration of neurosecretory protein VGF in tau aggregates that occurred specifically in neurofibrillary tangles in AD and in Pick bodies in PiD. VGF has been identified as interactor of wild-type and mutant forms of tau in recent proximity proteomic study [161], but has not been shown to be sequestered in phospho-tau lesions before. VGF co-aggregation with phospho-tau may play a crucial role in neurodegeneration. In physiological conditions, VGF is proteolytically processed in neuronal Gogi apparatus and secretory vesicles to produce active VGF-derived peptides that are then secreted by exocytosis to stimulate responses in neighboring cells [128]. Functions of these peptides include, but not limited to synaptogenesis, formation of long-term memory in hippocampus, and activation of microglia [10, 19, 89]. While it is hard to draw a definitive conclusion from our proteomic data on which of the VGF proteoforms is sequestered in phospho-tau lesions, peptides identified in our MS data indicate that it is most likely VGF precursor since some of them are outside of the boundaries of known VGF-derived peptides. Antibodies used in the immunofluorescent analysis were raised against central part of the VGF precursor molecule that contain only a few known active peptides, also indicating that their most likely target is VGF precursor molecule [communication with vendor]. If this is the case, sequestration of the VGF precursor in tau lesions likely results in disruption of active VGF-derived peptide production and secretion. In support of this, reduction of VGF peptides in CSF and brain has been reportedly found in different neurodegenerative conditions, including AD and FTD [6, 22, 43, 58, 59, 92, 93, 122, 131, 135, 140, 163]. While the mechanism that leads to reduction of VGF-derived peptides in these conditions is not known, it can be speculated that this effect may at least partially result from insufficient secretion of VGF-derived active peptides, since VGF-derived peptide TLQP-62 has been shown to stimulate VGF translation through VGF/BDNF/TrkB or mTOR/GPCR-dependent autoregulatory feedback loops [67, 90]. Interestingly, no change in VGF protein level was detected in PSP, along with no association between VGF and PSP phospho-tau lesions found in our study, again suggesting a link between VGF sequestration in aggregates and level of VGF protein and active peptides in brain [6].

VGF is increasingly recognized as a major player in neurodegenerative diseases. Most notably, VGF was recently identified as a key driver of AD by multiscale casual network analysis of multiomics data generated as part of the Accelerating Medicines Partnership-AD (AMP-AD) program from a large cohort of late-onset AD and control individuals [9]. The importance of VGF for AD pathology is supported by the findings that VGF partially mediates the effect of AD polygenic risk score on cognitive decline with the effect size comparable only to beta-amyloid and tau [156]. The most recent study established a link between higher *VGF* expression and slower cognitive decline and lower amount of neuropathologies in older adults [175]. This makes VGF not only a promising biomarker for neurodegenerative diseases, but also a perspective target for therapy. It has already been shown that increase of VGF level by VGF overexpression or upregulation of VGF translation rescued or partially rescued number of phenotypes in 5xFAD mouse model of AD [9]. Administration of VGF-derived peptide TLQP-21 in the same mouse model produced similar results, although in this case the effect was sex-specific and restricted to males [45]. Our results suggest that similar therapeutic approach can be also extended to certain tauopathies, particularly Pick’s disease. Although there are no good established animal models for 3R tauopathies, it would be interesting to test the effect of different modes of VGF administration in available tauopathy models, such as PS19 mouse line.

Our proteomics analysis also uncovered differential association of ferritin light chain protein FTL with phospho-tau aggregates in different tauopathies. We found that FTL is more strongly associated with AT8 phospho-tau in CBD than in AD, PiD, or PSP. Immunohistochemical analysis revealed that the most FTL positive tau lesion is astrocytic plaque that is a characteristic tau lesion in CBD. We also found co-localization of phospho-tau with FTL in tufted astrocytes in CBD and coiled bodies in CBD and PSP. In agreement with our findings, FTL was recently identified as phospho-tau interactor in brains from PSP patients [129]. We also found FTL co-localized with small phospho-tau glial lesions in PiD. No co-localization was detected in neuronal lesions (neurofibrillary tangles, globose tangles, Pick bodies, neuritic plaques and neuropil threads). This suggests that co-localization with FTL is a feature of glial tau pathology, in agreement with the evidence that ferritin light chain subunit is mostly expressed by glial cells, while neurons mostly express ferritin heavy chain subunit [24].

Our finding of higher FTL association with astrocytic plaques is in good agreement with the previous work that found higher level of FTL in CBD compared to PSP in caudate nucleus [44]. This work also showed co-localization of FTL with astrocytic plaques and tufted astrocytes. We now extend this finding to frontal cortex and show that association of FTL with phospho-tau is higher in CBD not only compared to PSP, but also to PiD and AD. However, in the study of Ebrahim et al., the increase of FTL in the caudate nucleus in CBD has been attributed to increase in FTL production by astrocytes [44]. While we cannot exclude that upregulation of FTL in astrocytes indeed occurs in CBD, we show that the majority of FTL co-localized with astrocytic plaques is associated with IBA1 positive microglia rather than with astrocytes. The difference between our results may arise from regional differences between frontal cortex and caudate nucleus, but also from the fact that different microglial markers have been used for co-immunostaining of microglia with FTL. In the study of Ebrahim et al. HLA-DR was used as a microglial marker [44]. However, it has been shown before that HLA-DR and FTL-positive microglia have distinct morphological appearance, suggesting that they may represent distinct microglial subpopulations [94]. In this study we used IBA1 that has been shown to be expressed in high level in microglia subpopulations with high expression of FTL [74].

Our study demonstrates that infiltration of astrocytic plaques with FTL-positive microglia is a characteristic feature of CBD pathology. High FTL expression in microglia has been previously suggested to be a marker of dystrophic microglial phenotype [94]. It is corroborated by our observations, as microglial cells associated with astrocytic plaque pathology in CBD have morphological appearance of dystrophic microglia. It is not entirely clear what exactly causes increased microglial FTL expression and dystrophic phenotype. One possibility is imbalance in iron metabolism, as it has been shown that FTL-positive microglia have higher iron-load than other microglial subtypes. We did not evaluate iron load in our CBD cases in this study, but in previous report staining with Prussian blue did not detect differences in iron content that could explain changes in FTL expression between CBD and PSP [44]. It has also been hypothesized that dystrophic microglial phenotype can be a caused by microglia exhaustion caused by unsuccessful attempts to phagocyte insoluble protein aggregates [153]. In any case, the increase in proportion of FTL-positive microglia has been associated with neurodegeneration and is likely to play an important role in pathology progression [145]. It will be important to assess the role of FTL-positive microglia in CBD pathology in future studies.

## Acknowledgements

We thank the patients and their families for participating in and contributing to these research studies and their generous donation of brain samples for research. This work was supported by NIH RF1AG076122 to WR, RF1AG076122-S1 supplement to AM, and a Chan-Zuckerberg Initiative (CZI) Collaborative Pairs Pilot Project Award to WR and MEM.

---

**Supplementary Fig. 1:**
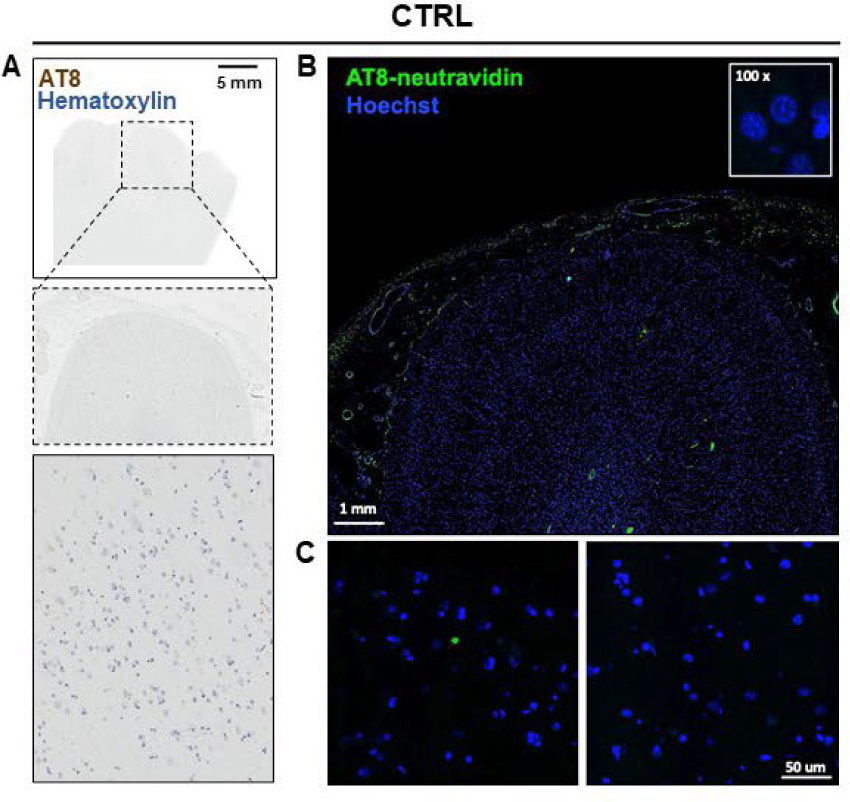
AT8 staining in frontal cortex from cognitively unaffected individual. (**A**) Control frontal cortex immunostained with AT8 phospho tau antibody and developed with DAB. (**B**) Proximity biotinylation using AT8 ProPPr.

**Supplementary Fig. 2.**
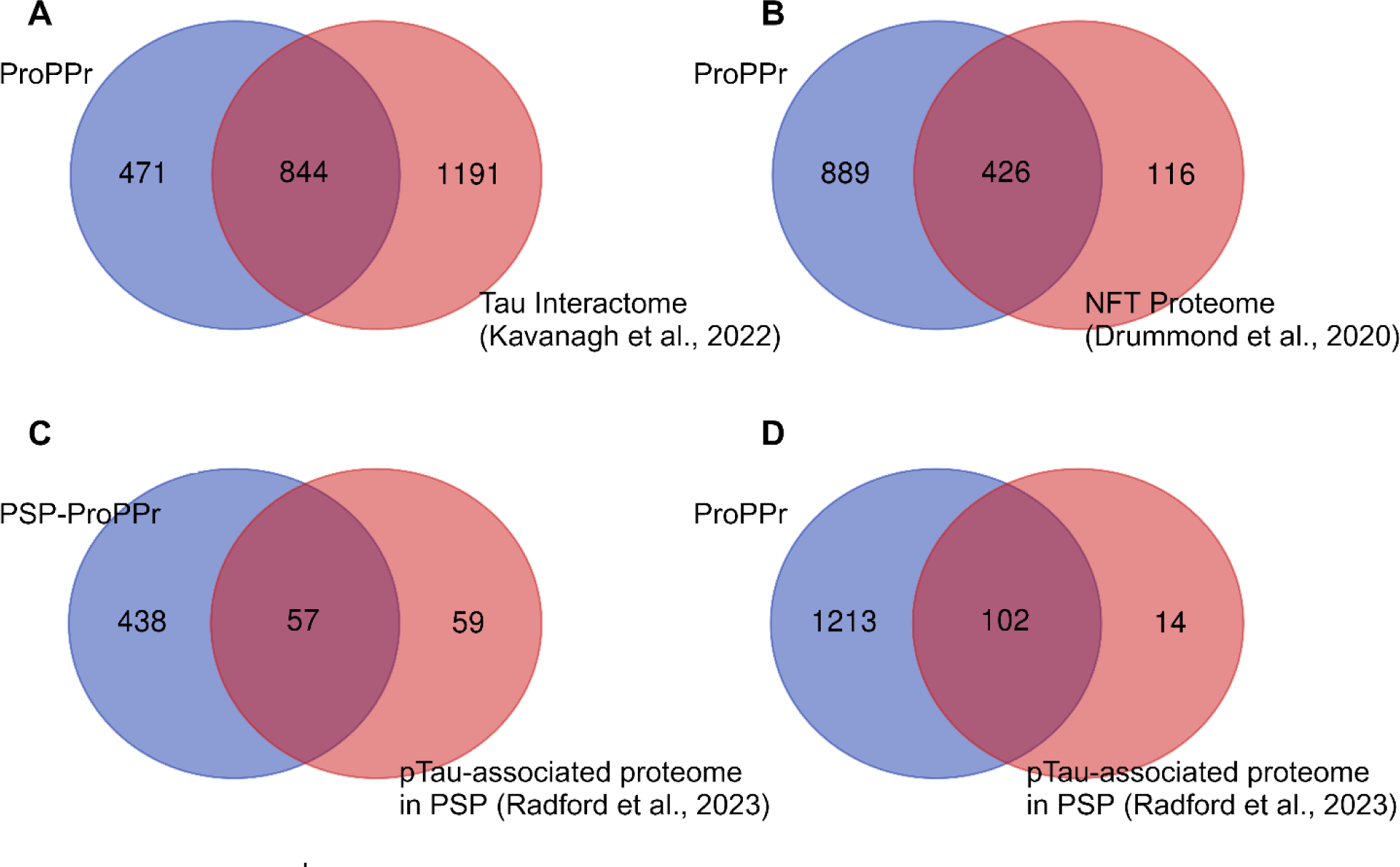
Comparison of the obtained set of phospho-tau associated proteins with previously published data. **(A)** Venn Diagram for the overlap between our set of phospho-tau-associated proteins and tau interactors reported in previous studies [73]. **(B)** Venn Diagram for the overlap between our set of phospho-tau-associated proteins and the proteome of neurofibrillary tangles in AD [41]. **(C)** Venn Diagram for the overlap between our set of phospho-tau-associated proteins in PSP and the PSP-associated proteome examined by BAR-MS method [129]. **(D)** Venn Diagram for the overlap between our set of phospho-tau-associated proteins found across the tauopathies and the PSP-associated proteome examined by BAR-MS method [129].

## Notes

### Competing Interest Statement

The authors have declared no competing interest.

